# SOD1 Mediates Lysosome-to-Mitochondria Communication and its Dysregulation by Amyloid-β Oligomers

**DOI:** 10.1101/2021.06.14.448281

**Authors:** Andrés Norambuena, Xuehan Sun, Horst Wallrabe, Ruofan Cao, Naidi Sun, Evelyn Pardo, Nutan Shivange, Dora Bigler Wang, Lisa A. Post, Heather A. Ferris, Song Hu, Ammasi Periasamy, George S. Bloom

## Abstract

Altered mitochondrial DNA (mtDNA) occurs in neurodegenerative disorders like Alzheimer’s disease (AD); how mtDNA synthesis is linked to neurodegeneration is poorly understood. We discovered <u>N</u>utrient-induced <u>M</u>itochondrial <u>A</u>ctivity (NiMA), an inter-organelle signaling pathway where nutrient-stimulated lysosomal mTORC1 activity regulates mtDNA replication in neurons by a mechanism sensitive to amyloid-β oligomers (AβOs), a primary factor in AD pathogenesis. Using 5-ethynyl-2’-deoxyuridine (EdU) incorporation into mtDNA of cultured neurons, along with photoacoustic and mitochondrial metabolic imaging of cultured neurons and mouse brains, we show these effects being mediated by mTORC1-catalyzed T40 phosphorylation of superoxide dismutase 1 (SOD1). Mechanistically, tau, another key factor in AD pathogenesis and other tauopathies, reduced the lysosomal content of the tuberous sclerosis complex (TSC), thereby increasing NiMA and suppressing SOD1 activity and mtDNA synthesis. AβOs inhibited these actions. Dysregulation of mtDNA synthesis was observed in fibroblasts derived from TS patients, who lack functional TSC and elevated SOD1 activity was also observed in human AD brain. Together, these findings imply that tau and SOD1 couple nutrient availability to mtDNA replication, linking mitochondrial dysfunction to AD.

## Introduction

The human brain represents ∼2% of the total body mass, but one fifth and one quarter of the total daily oxygen and glucose expenditure occurs there, respectively (Rolfe and Brown, 1997). Exquisite regulation of this costly oxygen and glucose budget is crucial to cognitive functions, especially in humans. The complexity of this daunting task multiplies when considering metabolic adjustments needed to support the activity of specific neuronal populations. It is not surprising, therefore, that much remains to be learned about how neurons orchestrate signaling pathways that control mitochondrial activity.

Reduction in brain glucose utilization is a well-known feature in cognitive disorders like Alzheimer’s disease (AD) (Camandola and Mattson, 2017). Fuoro-2-deoxyglucose positron-emission tomography (FDG-PET) studies of human brains have shown a larger decline in glucose utilization in the hippocampus and cortex in AD brain as compared to individuals without dementia (Kapogiannis and Mattson, 2011; Crane *et al*., 2013; Croteau *et al*., 2018; Gordon *et al*., 2018). At the organ level, PET studies tracking oxygen-15 showed that cerebral metabolic rate of oxygen is reduced in several AD brain regions and its decline positively correlates with severity of dementia (Ishii *et al*., 1996; Collins *et al*., 2015). As the sole oxygen-consuming organelle, mitochondria not only are the main cellular source of nutrient-derived chemical energy in the form of ATP, but also are responsible for producing ∼90% of the total cellular ROS (Balaban, Nemoto and Finkel, 2005), which *per se* serve as physiological envoys as well as noxious molecules when in excess (Shadel and Horvath, 2015). Along these lines, mitochondrial dysfunction has been long linked to a variety of neurodegenerative disorders (Chen and Chan, 2009; Sheng and Cai, 2012; Burté *et al*., 2014), in particular AD (Calkins *et al*., 2011; Manczak, Calkins and Reddy, 2011; DuBoff, Feany and Götz, 2013). At the molecular level, deficient brain energy metabolism has been linked to both compromised expression of electron transport chain proteins (Cottrell *et al*., 2001; Valla, Berndt and Gonzalez-Lima, 2001; Adav, Park and Sze, 2019) and mtDNA mutations in neurons (Mosconi *et al*., 2007). These observations not only highlight a maternally inherited risk factor in AD (Mosconi *et al*., 2007; Liang *et al*., 2008), but also suggest that dysregulation of mechanisms regulating the mitochondrial genome may account for brain energy metabolism deficiencies.

Mitochondria play a critical role in cell and body physiology by regulating the homeostasis of lipids, calcium and reactive oxygen species (ROS), cell death, inflammasome activation and others (Nunnari and Suomalainen, 2012; Labbé, Murley and Nunnari, 2014; Zhong *et al*., 2018). Mitochondrial dynamics and function are highly regulated by extracellular cues or global changes to enable timely adjustment on mitochondrial metabolism. This process also involves the coordinated expression of >1000 nuclear-encoded genes and those encoded by mtDNA (Nunnari and Suomalainen, 2012; Suomalainen and Battersby, 2017). Neurodegenerative disorders (DuBoff, Feany and Götz, 2013; Burté *et al*., 2014), type 2 diabetes (De Felice and Ferreira, 2014) and cancer (Vyas, Zaganjor and Haigis, 2016), have all been linked to defects in mtDNA maintenance (Trifunovic *et al*., 2004; Nunnari and Suomalainen, 2012). Mitochondria are the only animal cell organelles that contain extra-genomic coding DNA. Of the 90 proteins that constitute the electron transport chain machinery in mammals, 13 are encoded by mtDNA (Gustafsson, Falkenberg and Larsson, 2016). No mutations in mtDNA have been directly linked to AD, but brain oxidative metabolism deficiencies are associated with mtDNA mutations that affect expression of energy metabolism genes in posterior cingulate neurons (Mosconi *et al*., 2007; Liang *et al*., 2008). Furthermore, diminished mitochondrial functioning in AD is associated with single nucleotide polymorphisms, germline variants, deletions affecting the expression of electron transport chain genes and reduced base-excision repair in mtDNA (Wang et al., 2020). Still, the mechanisms coordinating nutrient availability, cellular redox states and mtDNA maintenance are not fully understood.

Superoxide dismutases (SODs) are metallo-enzymes that catalyze removal of superoxide free radicals (Miao and St. Clair, 2009). Copper and Zinc SOD (CuZnSOD or SOD1), and Manganese SOD (MnSOD or SOD2) are conserved intracellular proteins (Valentine, Doucette and Potter, 2005); SOD1 is the major SOD that is widely distributed throughout the cytosol, mitochondrial intermembrane space and the nucleus, whereas SOD2 exclusively resides in mitochondria (Weisiger and Fridovich., 1973). SOD1 activity restricts the superoxide production during mitochondrial respiration, directly controlling cellular ROS levels and oxidative damage. In addition, recent evidence suggests that SOD1 can directly modulate respiration by stabilizing casein kinase 1, a mitochondrial respiration repressor (Reddi and Culotta, 2013). SOD1 dysfunctions are well-known to cause ALS (Valentine, Doucette and Potter, 2005) and seems to play a role in cancer (Che *et al*., 2016). Although the potential involvement of SOD1 in AD has been suggested before (Yoon *et al*., 2009; Murakami *et al*., 2011), mechanistic evidence remains elusive.

Aβ peptides and tau can work in coordination to disrupt neuronal function at different levels, including organelle functioning (Guo *et al*., 2017; Norambuena *et al*., 2018; Polanco *et al*., 2018). For example, we recently discovered an inter-organelle signaling pathway, NiMA (**N**utrient-**i**nduced **M**itochondrial **A**ctivity) (Norambuena *et al*., 2018), in which activation of the lysosome-associated multi-subunit kinase, mechanistic target of rapamycin complex 1 (mTORC1) by insulin or amino acids, leads to inhibition of mtDNA synthesis and increase mitochondrial oxidative phosphorylation (OXPHOS) *in vitro* and *in vivo* (Norambuena *et al*., 2018). In addition, our group previously found that soluble AβOs trigger lipid raft-and Rac1dependent mTORC1 activity at the plasma membrane (PM), by a mechanism dependent on tau (Norambuena *et al*., 2017). As a consequence, AβOs trigger neuronal cell cycle re-entry, a harbinger of neuron death in AD (Arendt *et al*., 2010), and inhibit NiMA as well (Norambuena *et al*., 2018). The molecular mechanism regulating the NiMA signaling pathway is still not fully understood, but the recent addition of SOD1 to the expanding list of proteins regulated by mTORC1(Tsang *et al*., 2018), opened the possibility that SOD1 is an upstream NiMA regulator and a mediator of its disruption by the AβO-tau axis in AD.

Using 5-ethyl-2’-deoxyuridine (EdU) incorporation into mtDNA, metabolic imaging at single-cell and tissue levels by 2-photon fluorescence lifetime microscopy (2P-FLIM) (Norambuena *et al*., 2018), along with multi-parametric photoacoustic microscopy (MP-PAM); which enables label-free, simultaneous high-resolution intravital imaging of the cerebrovascular parameters including oxygen saturation in hemoglobin (sO_2_) and cerebral blood flow (CBF) (Ning *et al*., 2015), we now show that tau-dependent activation of lysosomal mTORC1 by insulin inhibits SOD1 activity in neurons by a mechanism that requires phosphorylation of SOD1 at T40. This allows mTORC1 to downregulate mtDNA synthesis in the presence of nutrients, a process that we found to be dysregulated by AβOs. SOD1 activity was stimulated by AβOs in cultured neurons and was elevated in human AD brain. Finally, pharmacologically inhibiting SOD1 activity with ATN-224 restrained mtDNA synthesis and inhibited respiration in live cells and in the mouse cortex. Thus, SOD1 is a part of a neuronal nutrient-sensing machinery that is based on tau and mTORC1, and functionally connects extracellular cues to mitochondrial functioning, enabling nutrients to sustain OXPHOS and prevent unwanted mtDNA synthesis. It is possible that disruption of NiMA contributes to oxidative damage and propagation of mtDNA mutations linked to diseases such as AD (Blanchard *et al*., 1993; Corral-Debrinski *et al*., 1994; Chagnon *et al*., 1999; Carrieri *et al*., 2001; Coskun, Beal and Wallace, 2004; Van Der Walt *et al*., 2004; Lakatos *et al*., 2010; Hudson *et al*., 2012; Inczedy-Farkas *et al*., 2014; Chen *et al*., 2016), ALS (Wiedemann *et al*., 2002) and tuberous sclerosis (TS) (Sakamoto *et al*., 2018).

## Results

### Tau regulates lysosomal mTORC1 activity and mtDNA synthesis

To gain a better understanding of the molecular mechanism regulating mtDNA synthesis through the NiMA pathway, we first asked whether it is influenced by tau. We began by comparing the incorporation of 5-ethyl-2’-deoxyuridine (EdU) into mtDNA in mouse primary neuron cultures from WT or tau KO mice. As shown previously by our group (Norambuena *et al*., 2018), WT mouse neurons readily incorporate EdU into mitochondrial nucleoids in a 3 hour incubation period, mainly in neuronal perikarya. Under these conditions, an average of ∼16 active EdU foci per neuronal perikaryon were detected (Figure 1A and B). In agreement with our previous observations (Norambuena *et al*., 2018), we also observed that the number of actively replicating nucleoids was reduced by ∼30% when lysosomal mTORC1 activity was stimulated with a mixture of amino acids (arginine and leucine, or R+L) plus insulin (Figure 1A and B). By contrast, the number of nucleoids in tau KO mouse neurons was ∼2-fold greater than in WT neurons and was insensitive to nutrient stimulation (Figure 1A and B). Re-expression of full length 2N4R human tau in tau KO neurons reduced the basal levels of EdU uptake by ∼30% compared to uninfected neurons and restored EdU uptake sensitivity to nutrient stimulation (Figure 1C and D). These observations imply that tau expression promotes lysosomal mTORC1 activity, and by extension, suppresses mtDNA synthesis.

**Figure 1.**
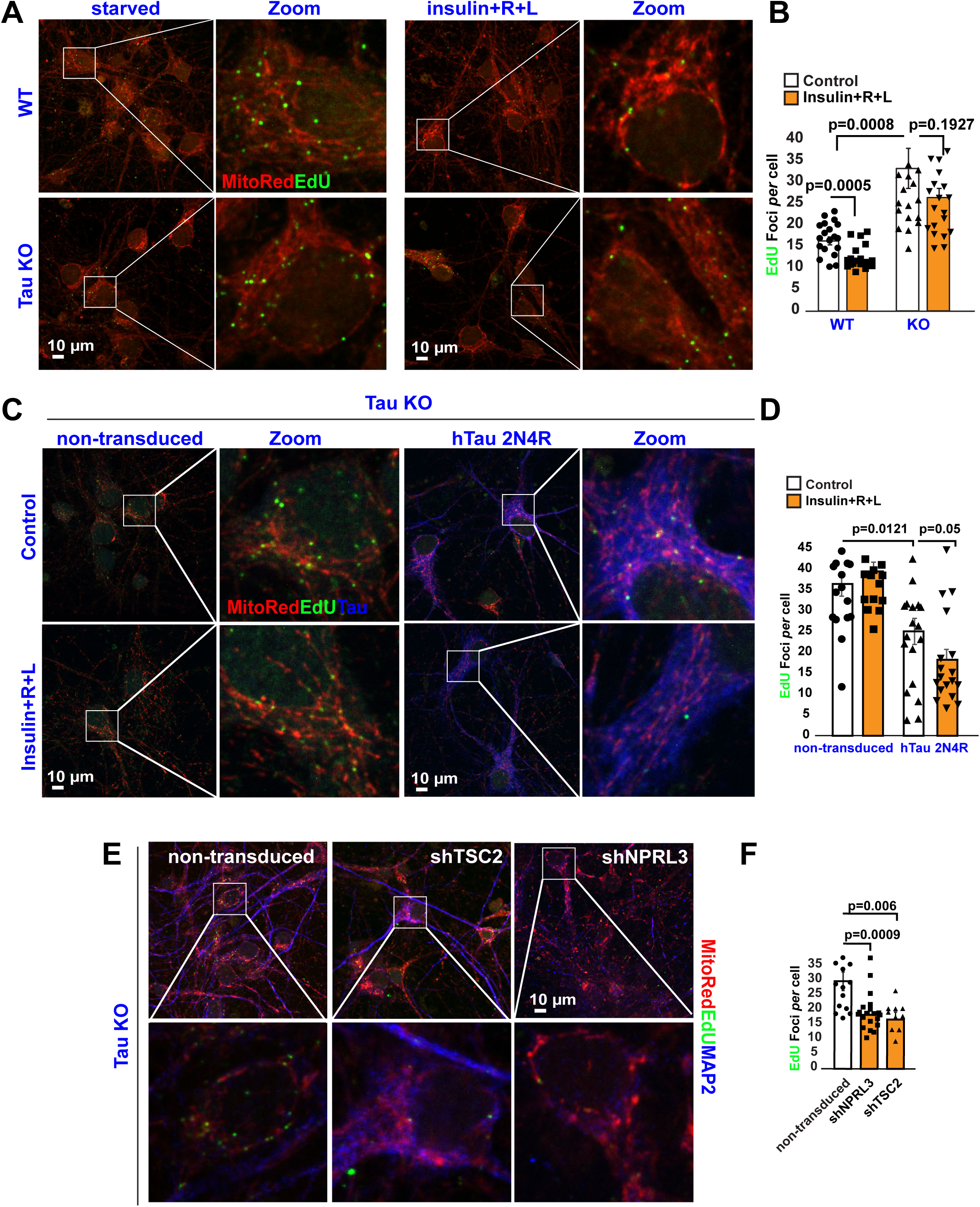
Tau regulates lysosomal mTORC1 signaling and mtDNA synthesis. **A-B** DNA replication in mitochondrial nucleoids was detected in both WT and tau KO mouse cortical neurons after a 3-hour pulse of the thymidine analog, EdU. Cells were labeled additionally with the mitochondrial marker, Mitotracker CMXRos (A). Quantification of EdU uptake into nucleoids showed that nutrients inhibit mtDNA replication (EdU foci/neurons) by ∼30% in WT (control: 16.35+/- 0.93; n=271 cells vs nutrients:11.9+/-0.68 EdU foci/cell; n=251 cells) but not in Tau KO neurons (control: 36.4+/- 2.9 EdU foci/cell; n=123 cells versus nutrients: 26.4+/-2.16 EdU foci/cell; n=120 cells). Each data point in the graph represents the average number of EdU foci/cell/field of view (FoV) recorded in this experiment and each FoV contained between 2-15 cells. Error bars represent ± s.e.m (B). Data are representative of at least three independent assays. **C-D** DNA replication in mitochondrial nucleoids was detected non-transduced tau KO neurons or after re-expressing human Tau 2N4R construct. Neurons were serum-starved in HBSS for 2 hours. Next, the cells were pulse-labeled for 3 hours with EdU (C). Note that forcing hTau expression alone yielded an ∼30% decrease in EdU incorporation into mtDNA in the in the absence of nutrients (36.48 +/-2.9 EdU foci/cell; n**=**123 cells vs 25.1 +/- 3.13 EdU foci/cell; n=75 cells), further decreasing by 20% in the presence of nutrients (control: 25.1 +/- 3.13 EdU foci/cell; n=75 cells vs nutrients: 18.27 +/- 2.5 EdU foci/cell; n=87 cells). Each data point in the graph represents the average number of EdU foci/cell/FoV recorded in this experiment and each FoV contained between 2-15 cells. Error bars represent ± s.e.m (D). Data are representative of at least three independent assays. **E-G** Genetic activation of mTORC1 activity in Tau KO neurons rescues insulin resistance effect on mtDNA synthesis. Neurons were serum-starved in HBSS for 2 hours. Next, the cells were pulse-labeled for 3 hours with EdU (Non-transduced: 29.8+/-3.13 EdU foci/cell; n=221 cells vs shNPRL3: 18.3 +/-1.24 EdU foci/cell; n=336 cells vs shTSC2: 16.8+/-2.5 EdU foci/cell; n=201 cells). Each data point in the graph represents the average number of EdU foci/cell/FoV recorded in this experiment and each FoV contained between 2-10 cells. Error bars represent ± s.e.m. Data are representative of two independent assays.

To test this idea directly, we monitored the effect of tau expression on lysosomal mTORC1 activity by expressing LysoTorcar, a biosensor of <u>lyso</u>somal m<u>TORC</u>1 <u>a</u>ctivity (Zhou *et al*., 2015). LysoTorcar corresponds to the full-length 4EBP1 protein, a well-known mTORC1 substrate for phosphorylation at T37, T46 and S65, flanked by Cerulean and Ypet fluorescent proteins. Phosphorylation of those sites can be monitored by western blotting and signifies mTORC1 activation, which can be reversed by dephosphorylation. Finally, the N-terminal ∼25% of LysoTorcar corresponds to LAMP1, which targets the fusion protein to lysosomes (Zhou *et al*., 2015) (Supplemental 1A). Consistent with its ability to report mTOR activity, basal LysoTorcar phosphorylation in HEK293 cells was reduced by ∼70% by the mTOR inhibitor, Torin 1 (Supplemental 1A). To test if tau affects mTORC1 activity, HEK293 cells were co-transfected with LysoTorcar, and either GFP or GFP-tau (human 0N4R isoform) for 24 hours. Then, cells were serum-starved for 1.5 hours before adding 100 nM insulin and R+L for another 25 minutes. Under baseline, unstimulated conditions, phosphorylation of LysoTorcar on T37 and T46 and S65 was ∼50% higher in cells expressing GFP-tau compared to GFP, and the level increased by an extra 50% when cells were stimulated with either insulin or R+L (Supplemental 1B). These results suggest that dysregulation of mtDNA synthesis in tau KO neurons occurs as a consequence of lower basal levels of mTORC1 activity. In fact, when the activity of lysosomal mTORC1 was genetically increased in tau KO mouse neurons by shRNA-mediated reduction of either of two suppressors of mTORC1 kinase activity, the TS complex (TSC) or GATOR1 complex (using shTSC2 or shNPRL3, respectively) we observed a ∼50% reduction in the number of replicating mitochondrial nucleoids (Figure 1E and F), reaching values similar to those detected in WT neurons (Figure 1A).

### Tau regulates lysosomal positioning of the TSC

Activation of lysosomal mTORC1 by nutrients can be achieved by R+L-stimulated recruitment of mTORC1 from cytosol to the lysosomal membrane (Saxton and Sabatini, 2017) or insulin-induced detachment from the lysosomal surface of the TSC complex (Menon *et al*., 2014). As shown by our group previously (Norambuena *et al*., 2017), stimulation of WT primary neuron cultures for 30 minutes with a mixture of R+L and insulin increased the amount of mTOR on the lysosomal surface, as judged by an increase in its co-localization with the lysosomal marker, LAMP1 (Figure 2A and C). A similar increase was also observed in tau KO neurons (Figure 2A and C). Interestingly, nutrient stimulation reduced the lysosomal content of TSC2, a subunit of the TSC complex, in WT neurons, but not in tau KO neurons (Figure 2B and D). Re-expressing 2N4R human tau in tau KO neurons (Western Blots in Figure 2G) did not alter mTOR co-localization with LAMP1 under nutrient-deprived conditions but trended towards enabling increased co-localization after nutrient stimulation (Figure 2E). On the other hand, tau re-expression in tau KO neurons led to reduced co-localization of TSC2 with LAMP1 in nutrient-deprived neurons, which decreased further after nutrient stimulation (Figure 2F). These collective results demonstrate that tau regulates lysosomal mTORC1 activity and mtDNA synthesis by regulating the lysosomal localization of the mTORC1 suppressor, TSC.

**Figure 2.**
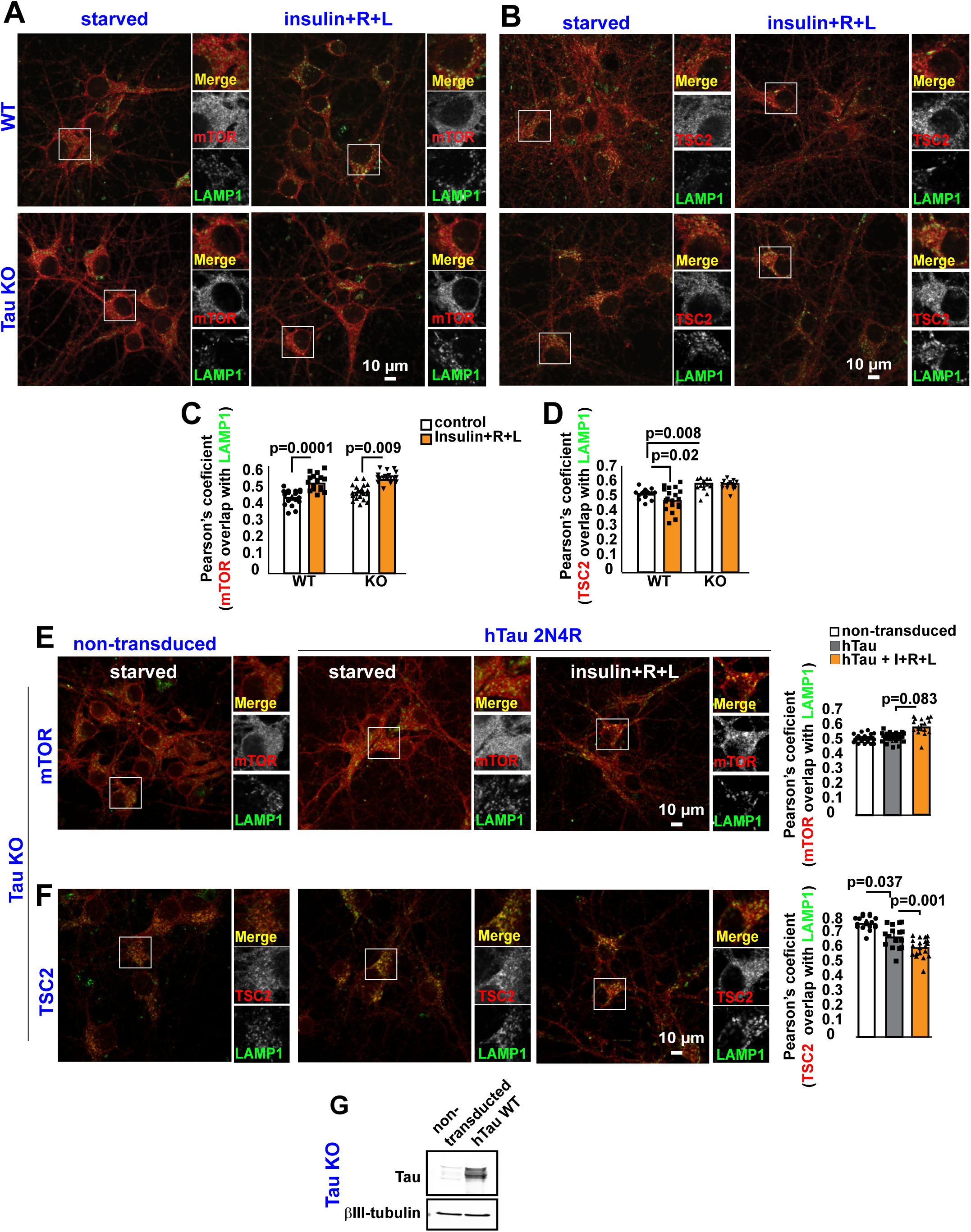
Tau Regulates TSC complex positioning on Lysosomes. **A-B** WT or Tau KO cortical neurons starved of insulin and amino acids for 2 hours were treated with insulin, or arginine and leucine (R+L) for 30 minutes and then were labeled by double immunofluorescence for mTOR and LAMP1, or TSC2 and LAMP1. Data are representative of at least three independent assays. **C-D** Quantification of experiments shown in A-B. Each data point in the graph represents the average Pearson’s coefficient/FoV recorded in this experiment and each FoV contained between 2-15 cells. Error bars represent ± s.e.m. Data are representative of at least three independent assays. **E-F** Non-transduced or human 2N4R Tau-expressing Tau KO cortical neurons starved of insulin and amino acids for 2 hours were treated with insulin, or R+L for 30 minutes, and then were labeled by double immunofluorescence for mTOR and LAMP1, or TSC2 and LAMP1. Each data point in the graph represents the average Pearson’s coefficient value/FoV recorded in this experiment and each FoV contained between 2-15 cells. Error bars represent ± s.e.m. Data are representative of at least three independent assays. **G** Representative western blot showing expression of human 2N4R Tau in Tau KO cortical neurons.

### mtDNA synthesis is insensitive to nutrients in TS human fibroblasts

TS is a developmental disorder caused by loss-of-function mutations in *TSC1* or *TSC2* whose protein products, TSC1 and TSC2, along with TBC1D7, constitute the TSC (Gao *et al*., 2002). TS patients develop benign, tuber-like tumors in brain, heart, kidney and skin, and are prone to cognitive deficits (Prather and de Vries, 2004), autism (Smalley, 1998), epilepsy (Thiele, 2004) and other neurological symptoms (Orlova and Crino, 2010). Since TSC is a major regulator of mTORC1 activity, we evaluated mtDNA synthesis in human fibroblasts derived from patients affected by TSC. Mutations in *TSC1* rendered mtDNA synthesis refractory to nutrient stimulation in human TSC fibroblasts (Figure 3A). Re-expressing WT human TSC1 in these cells not only increased the number of nucleoids in starved fibroblasts but also restored sensitivity to nutrient-stimulation (Figure 3B). These data reinforce our previous report (Norambuena *et al*., 2018) that malfunctioning of lysosomal mTORC1 causes mitochondria to be refractory to insulin and amino acids signaling in at least two diseases: in AD, in which AβOs triggers mTORC1 activation at the PM (Norambuena *et al*., 2017), and in tuberous sclerosis, due to loss of functional TSC complex.

**Figure 3.**
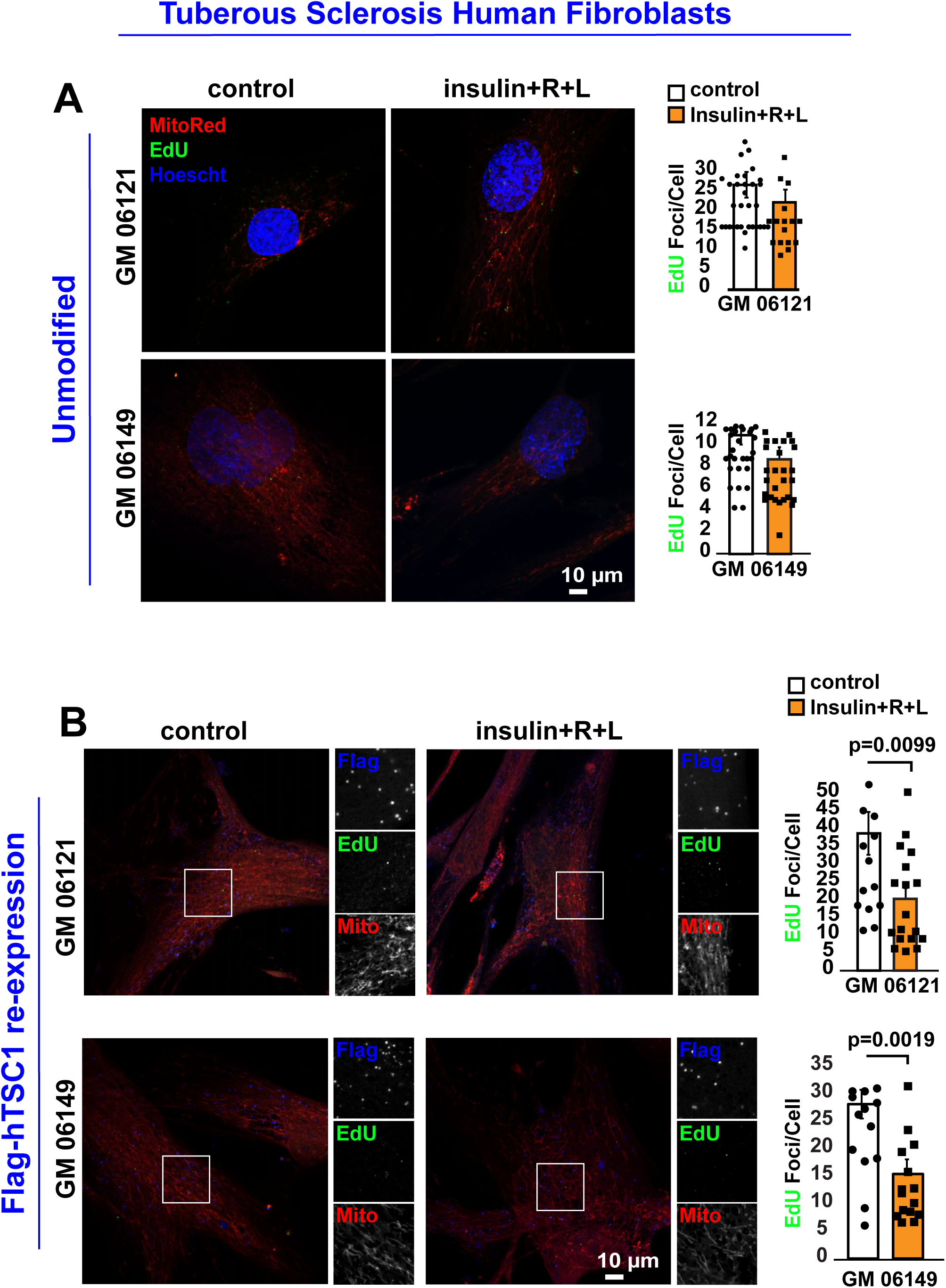
mtDNA synthesis is dysregulated in tuberous sclerosis fibroblasts. **A** DNA replication in mitochondrial nucleoids was detected in human fibroblasts from two patients harboring polymorphisms in TSC1 (cases# GM 06121 and GM 06149). Note that deficient TSC complex functioning in these fibroblasts renders mtDNA synthesis insensitive to regulation by nutrients. Each data point in the graph represents the average number of EdU foci/cell/FoV recorded in this experiment and each FoV contained between 2-5 fibroblasts. Error bars represent ± s.e.m. Data are representative of at least three independent assays. **B** Re-expression of human TSC1 rescues mtDNA synthesis response to nutrients. Note that expressing a functional TSC1 protein (likely restoring TSC complex normal functioning and mTORC1 activity) increased the number of EdU foci per fibroblast and rescued its sensitivity to nutrient stimulation. Expression of the Flag-WT TSC1 was corroborated by indirect immunofluorescence (IF) against the Flag epitope. Each data point in the graph represents the average number of EdU foci/cell/FoV recorded in this experiment and each FoV contained between 2-5 fibroblasts. Error bars represent ± s.e.m. Data are representative of at least three independent assays.

### Tau controls mtDNA synthesis through SOD1

mTORC1 phosphorylates and regulates the activity of numerous substrates (Hsu *et al*., 2011a; Yu *et al*., 2011). We previously showed that NiMA was independent of well-known mTORC1 substrates including S6K, 4EBP/eIF4E and BcL-xL (Norambuena *et al*., 2018). We did not, however, previously examine superoxide dismutase 1 (SOD1), which was recently shown to be an mTORC1 substrate (Tsang *et al*., 2018), and serves as the main regulator of the cytosolic redox state by consuming superoxide produced during mitochondrial respiration and generating the signaling molecule hydrogen peroxide (Veal, Day and Morgan, 2007). SOD1 thus emerged as a potential coordinator of extracellular cues and redox regulation with mitochondrial function through the NiMA pathway. To test whether SOD1 activity is part of the tau-mTORC1 signaling pathway, we first evaluated SOD1 activity in WT and tau KO neurons that were starved, or stimulated with insulin. SOD1 activity in WT neurons was reduced by ∼60% after 30 minutes of treatment with nutrients (Supplemental 2A). In contrast, SOD1 activity in tau KO neurons was reduced by insulin stimulation by only∼10% (Supplemental 2B). Conversely, re-expressing 0N4R human tau in tau KO mouse neurons decreased SOD1 activity by 40% in the absence of mTORC1 stimulation and decreased even more after nutrient treatment (Supplemental 2C). These collective results imply that tau regulates SOD1 through its regulation of lysosomal mTORC1 activity, and by extension SOD1 regulates mtDNA synthesis. Consistent with that interpretation, the basal level of EdU-positive mitochondrial nucleoids in neurons was reduced by ∼50% by the pharmacological SOD1 inhibitor, ATN 224, and by ∼75% by shRNA-mediated reduction of SOD1 (Figure 4A-D).

mTORC1 phosphorylates human SOD1 at T40 (Tsang *et al*., 2018), thereby downregulating SOD1 activity. To test whether phosphorylation of SOD1 on T40 was required for nutrient-mediated regulation of mtDNA synthesis we introduced single amino acid substitutions into a GFP-tagged WT human SOD1 fusion protein to produce phospho-mimetic (T40E) and phospho-null (T40A) mutants. Nutrient stimulation by insulin and R+L reduced the number of replicating nucleoids by 25% in mouse neurons expressing GFP-SOD1^WT^ (Figure 4E and F). In contrast, the number of nucleoids in neurons expressing GFP-SOD1^T40A^ under starvation conditions was ∼25% higher than in neurons expressing GFP-SOD1^WT^ and was insensitive to nutrient stimulation (Figure 4E and F). Conversely, expression of GFP-SOD1^T40E^ reduced the base-line number of nucleoids by 24% compared to expression of GFP-SOD1^WT^, reaching levels similar to cells expressing GFP-SOD1^WT^ after nutrient stimulation (Figure 4E and F). In addition, the number of mitochondrial nucleoids in GFP-SOD1^T40E^ expressing neurons was insensitive to nutrient stimulation (Figure 4E and F). Finally, the number of nucleoids in tau KO neurons, which showed lower levels of mTORC1 activity and double the number of EdU foci per cell compared to WT neurons, was decreased by ∼50% by expressing GFP-SOD1^T40E^ (Supplemental 3). Thus, phosphorylation at T40 in SOD1 regulates mtDNA synthesis through the tau-mTORC1 signaling pathway.

**Figure 4.**
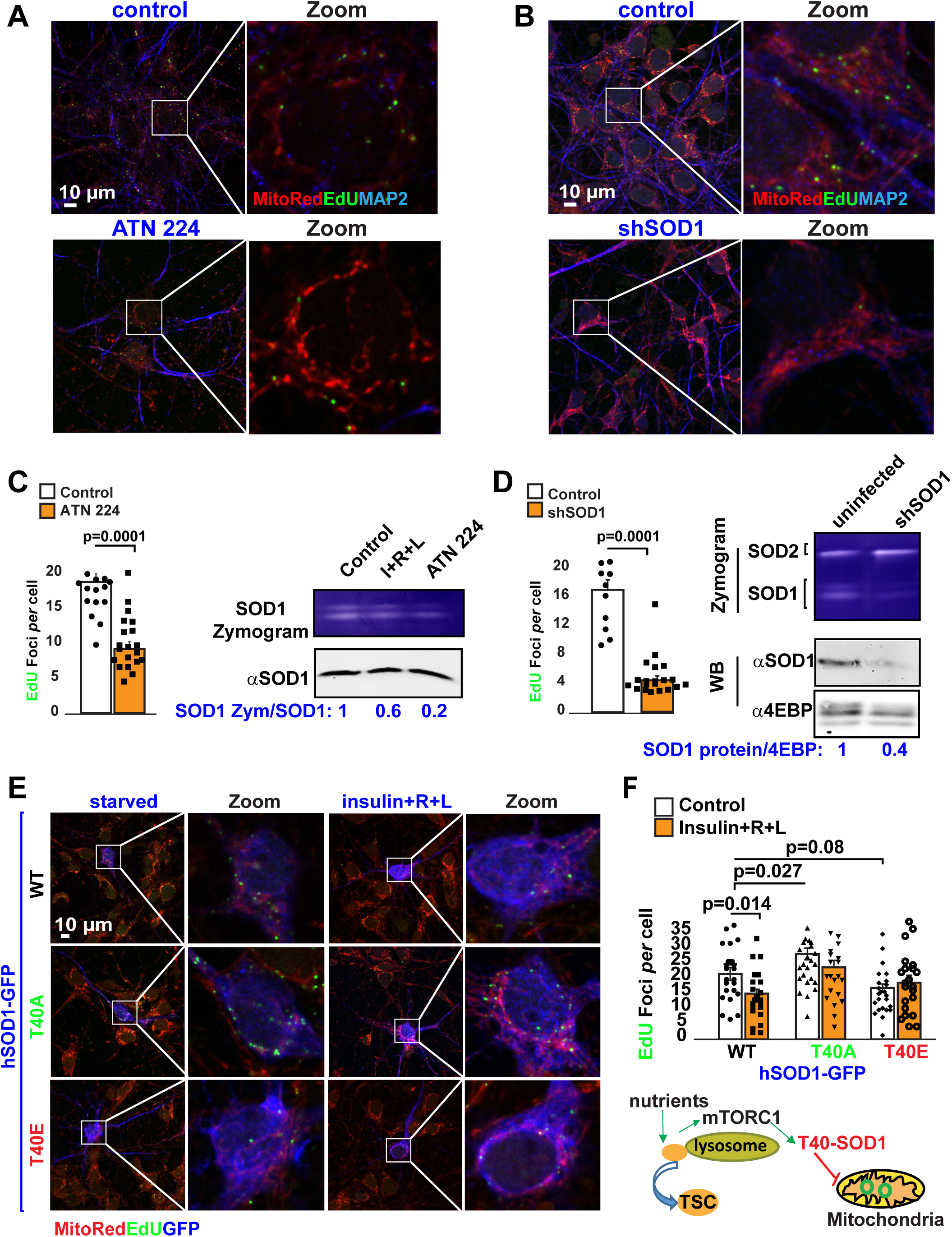
Threonine 40 on SOD1 regulates mtDNA synthesis through the NiMA pathway. **A-D** DNA replication in mitochondrial nucleoids was detected in WT mouse cortical neurons treated with the SOD1 inhibitor, ATN-224 (A), or in SOD1 knockdown neurons (B). Neurons were serum-starved in HBSS for 2 hours. Next, the cells were pulse-labeled for 3 hours with EdU. Quantification of EdU uptake into nucleoids showed that ATN-224 and SOD1 reduction inhibit mtDNA replication by ∼60% (control: 18.7±1.16 EdU foci/cell, n=332 vs ATN224: 9.1±1 EdU foci/cell, n=150 cells) and ∼70% (control: 16.9±1.4 EdU foci/cell, n=345 vs shSOD1: 4.4±0.6 EdU foci/cell, n=262), respectively. SOD1 activity under these conditions was evaluated using in-gel activity assays. Each data point in the graph represents the average number of EdU foci/cell/FoV recorded in this experiment and each FoV contained between 2-15 cells. Error bars represent ± s.e.m. Data are representative of three independent assays. **E** DNA replication in mitochondrial nucleoids was evaluated in WT mouse cortical neurons expressing hSOD1-GFP before and after nutrient stimulation. Quantification of EdU uptake into nucleoids showed that cells expressing either hSOD1^T40A^-GFP or hSOD1^T40E^-GFP renders mtDNA synthesis insensitive to nutrient regulation (hSOD1-GFP^WT^ control: 21.4+/-2.2 EdU foci/cell vs nutrients: 15+/-1.4 EdU foci/cell; vs hSOD1-GFP^T40E^ control:16.8+/-1.3 EdU foci/cell vs nutrients: 18.5+/-2.4 EdU foci/cell; vs hSOD1-GFP^T40A^ control: 28+/-1.9 EdU foci/cell vs nutrients: 24+/- 2.3 EdU foci/cell. Each data point in the graph represent the average number of EdU foci/GFP-expressing cell/FoV recorded in this experiment and each FoV contained between 1-2 cells. Error bars represent ± s.e.m. Data are representative of three independent assays.

### ATN 224 inhibits mitochondrial respiration in cultured cells and mouse brain

Having demonstrated that the tau-mTORC1-SOD1 signaling pathway regulates mtDNA synthesis, we then explored whether SOD1 activity is required for respiration in cultured HEK293 cells, mouse neurons, or in mouse brain. For the cultured cells we used 2-photon fluorescence lifetime imaging (2P-FLIM) to monitor NAD(P)H lifetimes and the degree to which these co-enzymes associate with partner enzymes in perinuclear mitochondria, as shown previously (Norambuena *et al*., 2018). The “bound fraction” of NAD(P)H, expressed as “a_2_%” (Lakowicz, 2006), positively correlates with mitochondrial respiration (Norambuena *et al*., 2018) and biosynthetic pathways (Blacker *et al*., 2014). Following serum starvation in Hank’s balanced salt solution (HBSS) for 2 hours, individual fields of view were assayed by 2P-FLIM before and 60 minutes after stimulation of the cells with Bis(choline)tetrathiomolybdate (ATN-224), a copper chelator and SOD1 inhibitor (Chidambaram, Barnes and Frieden, 1984). ATN-224 caused a similar decrease in enzyme-bound NAD(P)H in both HEK293 and neurons, signifying a downregulation in oxidative phosphorylation (OXPHOS) activity (Figure 5A and B). In addition, ATN-224 significantly reduced oxygen consumption in neuron cultures (Figure 5C).

**Figure 5.**
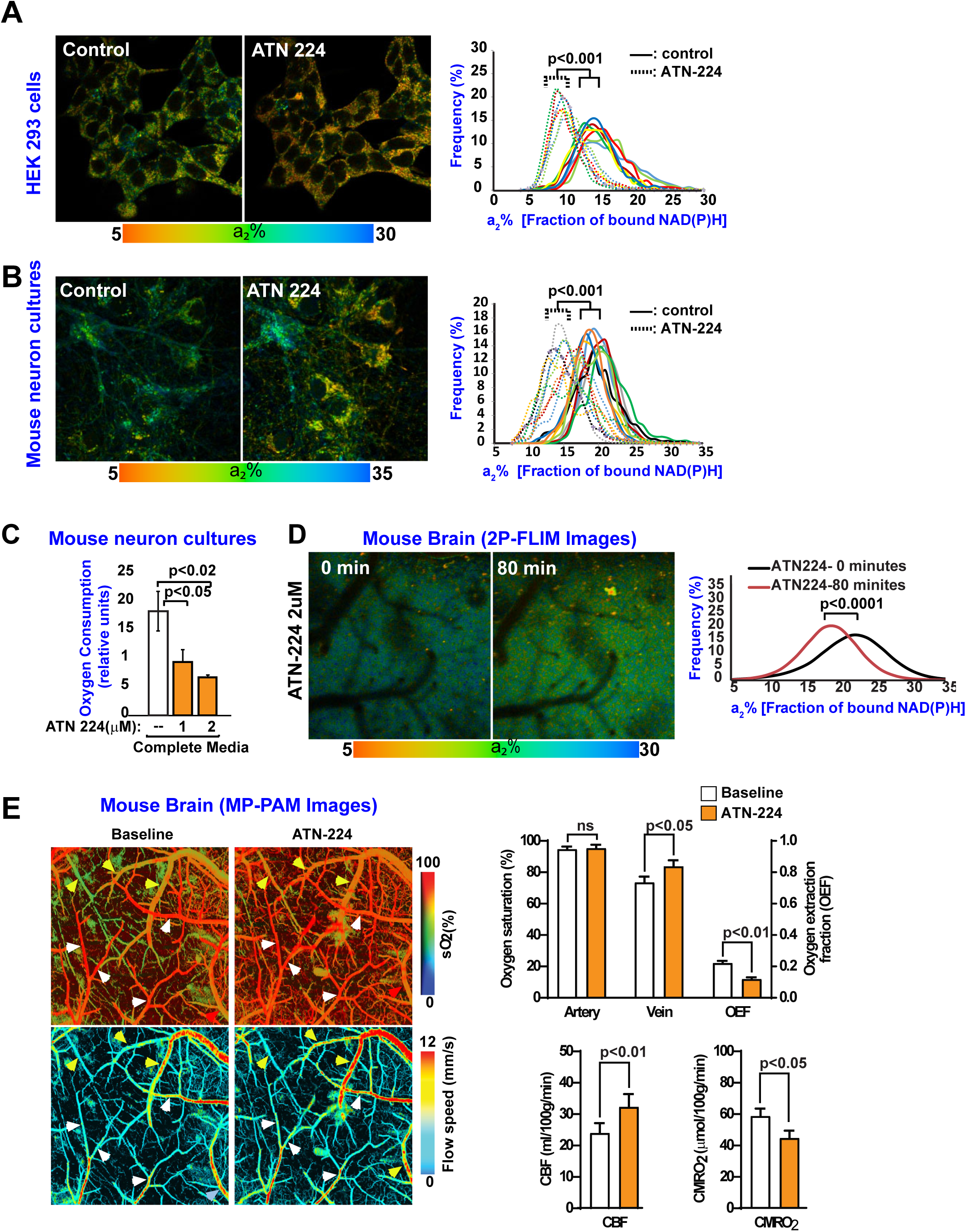
ATN-224 regulates mitochondrial activity in live cells and brain. **A-B** a2% values (fraction of bound NAD(P)H; baseline control, and 80 minutes after ATN-224 treatment) were recorded pixel by pixel for HEK293 cells (A) and mouse neurons (B). Each colored line in the panel graphs refers to a single field of view containing ∼15 cells, and each pair of solid and dotted lines (same color) refers to the same field of view before and after ATN-224 treatment, respectively. All statistical analyses were performed using Student’s two-tailed unpaired t-tests. Data are representative of two independent assays. **C-D** ATN-224 reduces oxygen consumption in mouse neurons in culture and live mouse brain. Mouse neurons cultures were kept in complete medium in the absence or presence of the SOD1 inhibitor, ATN-224. One hour later, oxygen consumption was measured. The data were collected from three independent assays in which each experimental condition contained six replicates. Error bars represent ± s.e.m (C). 2P-FLIM images (a2% values) of the live mouse brain cortex were recorded pixel by pixel before and after an 80 min treatment with ATN-224 through an open skull window. Statistical analyses were performed using Student’s two-tailed unpaired t-test. Data are representative of three independent assays. **E** MP-PAM imaging of wild-type (WT) mouse cerebral cortex through an open-skull window 80 min. after topical application of ATN-224. An increase in blood oxygenation of the cortical vasculature was observed, indicating decreased oxygen extraction and consumption due to downregulation of mitochondrial activity. Data were obtained from three mice, each of which was measured once for O_2_ saturation, oxygen extraction fraction (OEF), and cerebral metabolic rate of oxygen (CMRO2). Error bars represent ± s.e.m. White arrows: Arteries; yellow arrows: Veins.

We next asked whether SOD1 activity also regulates mitochondrial respiration at the tissue level *in vivo*. First, we used 2P-FLIM to monitor changes in NAD(P)H lifetimes in live mouse brain. This approach revealed that topical application of ATN-224 for 80 minutes through a cranial window caused a substantial decrease in the fraction of enzyme-bound NAD(P)H (Figure 5D). In parallel, we applied multiparametric photoacoustic microscopy (MP-PAM) to measure oxygen-metabolic responses in the live mouse brain (Cao *et al*., 2017). MP-PAM imaging through a cranial window enables simultaneous, high-resolution imaging of total hemoglobin concentration (C_Hb_), oxygen saturation of hemoglobin (_S_O2), and cerebral blood flow (Ning *et al*., 2015). This approach revealed that ATN-224 caused significant increases in venous _S_O2, without changes in cerebral blood flow (CBF; Figure 5E). These hemodynamic responses to ATN-224 were accompanied by decreases in both oxygen extraction fraction (OEF) and the cerebral metabolic rate of oxygen (CMRO_2_; Figure 5E). Altogether, these results suggest that mTORC1-mediated inhibition of SOD1, through phosphorylation of T40, inhibits mtDNA synthesis and stimulates mitochondrial respiration in both proliferating and non-proliferating cells. The inhibitory effect of ATN-224 on OXPHOS in live cells and mouse brain seems counterintuitive, but likely occurs as a consequence of inhibiting cytochrome c oxidase through copper sequestration (Nair and Mason, 1966), rather than by SOD1 inhibition.

### Tau controls nutrient-mediated suppression of mtDNA synthesis and its dysregulation by AβOs

We previously reported that AβO-mediated disruption of NiMA requires tau expression (Norambuena *et al*., 2018). As tau expression in tau KO neurons restores normal lysosomal mTORC1 signaling and mtDNA synthesis in neurons (Figure 1), we tested whether tau is also required for nutrient-induced inhibition of mtDNA synthesis, and the upregulation of mtDNA synthesis triggered by AβOs (Norambuena *et al*., 2018). Contrary to the inhibitory action of nutrients on mtDNA synthesis observed in WT mouse neurons (Figure 1A), nutrients did not significantly decrease the nucleoid content of tau KO neurons (Figure 6A). Likewise, the number of nucleoids in tau KO neurons were not affected by AβOs in either the absence or presence of nutrients (Figure 6A). Tau re-expression not only reduced the basal level of nucleoids by 30%, but also restored inhibition of mtDNA synthesis by nutrient stimulation (Figure 6B). Furthermore, tau re-expression in tau KO neurons re-established the ability of AβOs to increase mtDNA synthesis (Figure 6B) and to abolish the inhibitory effect of nutrients (Figure 6B). Having established that tau permits proper lysosomal mTORC1 signaling and its counteraction by AβOs, we then asked whether T40 phosphorylation on SOD1 mediates AβO dysregulation of mtDNA synthesis. AβO-treatment increased the number of nucleoids by ∼30% in neurons expressing either GFP or GFP-SOD1^WT^, but not in GFP-SOD1^T40E^ expressing neurons (Figure 6C). AβOs therefore signal through the tau-mTORC1-SOD1 signaling pathway to dysregulate mtDNA synthesis.

**Figure 6.**
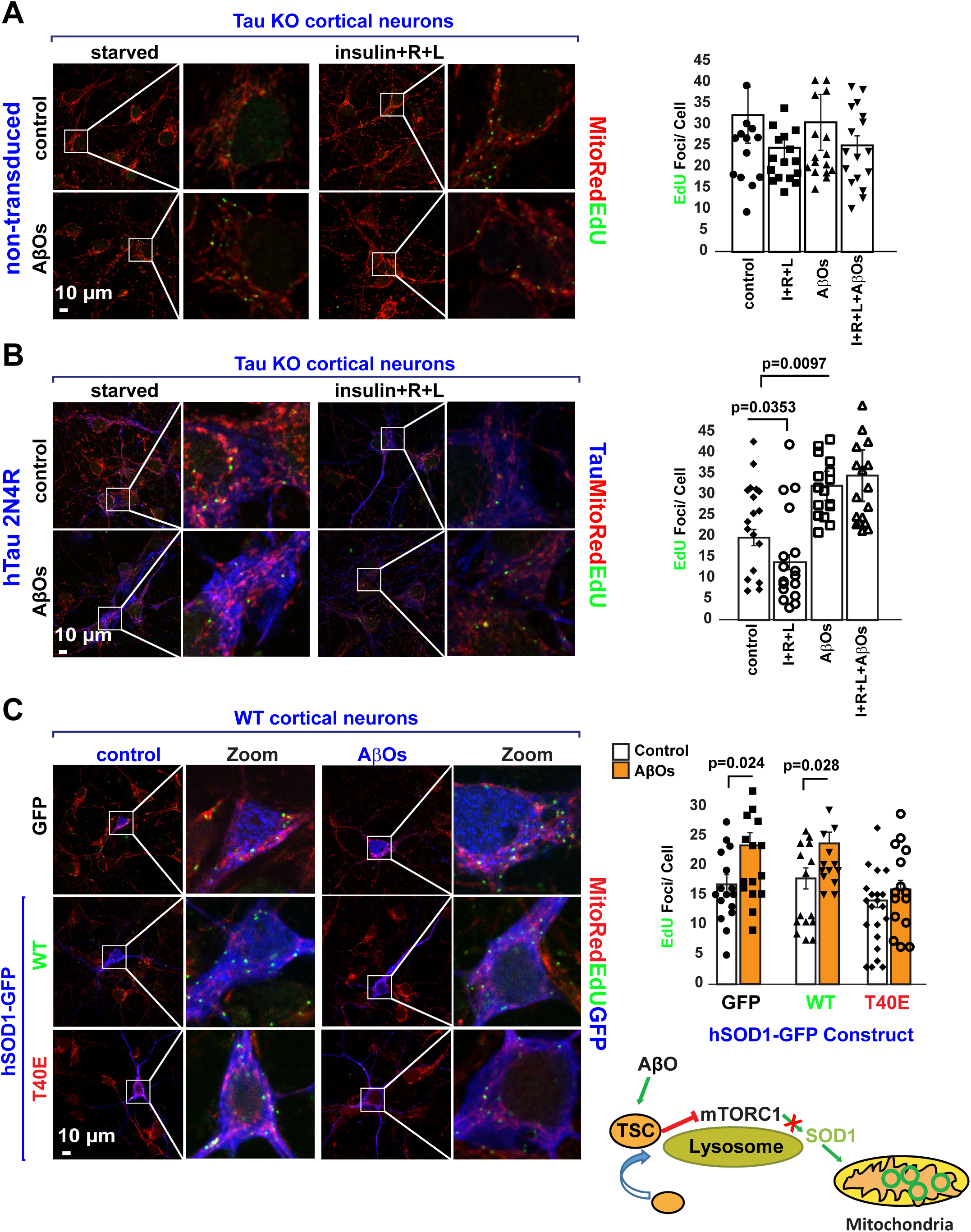
AβO-mediated dysregulation of mtDNA synthesis requires tau and is mediated by SOD1 T40. **A** DNA replication in mitochondrial nucleoids was detected in tau KO mouse neurons. Neurons were serum-starved in HBSS for 2 hours. Next, the cells were pulse-labeled for 3 hours with EdU in the absence or presence of nutrients (a mixture of insulin and R+L, AβOs, or a combination of both. Quantification of EdU uptake into nucleoids showed that mtDNA replication was insensitive to either nutrient stimulation (control: 32± 6.7 EdU foci/cell, n=80 vs nutrients: 25± 2.6 EdU foci/cell; n=102), AβO (31±6.6 EdU foci/cell, n=102) or AβOs plus nutrients (25.2±2.2 EdU foci/cell, n=71). Each data point in the graph represents the average number of EdU foci/cell/FoV recorded in this experiment and each FoV contained between 1-2 cells. Error bars represent ± s.e.m. Data are representative of three independent assays. **B** Cortical mouse neurons from tau KO mice were transduced with lentivirus to express hTau 2N4R, which not only rescued the ability of tau KO neurons to reduce mtDNA synthesis in the presence of nutrients (control: 19.8±1.9 EdU foci/cell n=69 cells vs nutrients: 13.9±1.8 EdU foci/cell n=65 cells) but also to increase it upon AβO treatment (control:19.8±1.9 EdU foci/cell, n=69 cells vs AβO: 32±3.5 EdU foci/cell, n=55 cells vs AβO+nutrients: 34.6±6.2 EdU foci/cell n=55 cells). Each data point in the graph represent the average number of EdU foci/GFP-expressing cell/FoV recorded in this experiment and each FoV contained between 1-2 GFP-expressing cells. Error bars represent ± s.e.m. Data are representative of three independent assays. **C** DNA replication in mitochondrial nucleoids was detected in WT mouse cortical neurons expressing either GFP, hSOD1^WT^-GFP or hSOD1^T40E^-GFP. Note that AβO treatment increase mtDNA synthesis by ∼45% in both GFP (control:17±1.7 EdU foci/cell n=30 cells vs AβO: 23.5±2.2 EdU foci/cell, n=35 cells) and hSOD1^WT^-GFP (control: 18±1.8 EdU foci/cell n= 35 cells vs AβO: 24±1.9 EdU foci/cell, n=33 cells) but not in hSOD1^T40E^-GFP expressing neurons (control: 16.2±1.4 EdU foci/cell n= 40 cells vs AβO: 18.41±1.8 EdU foci/cell n=33 cells). Each data point in the graph represents the average number of EdU foci/ GFP-expressing cell/FoV recorded in this experiment and each FoV contained between 1-2 cells. Error bars represent ± s.e.m. Data are representative of three independent assays.

### SOD1 activity is elevated in Alzheimer’s disease human brain

Previous work by our group and others has shown that AβOs impair insulin/IGF1 signaling (De Felice and Ferreira, 2014; Norambuena *et al*., 2017). Mechanistically these effects are explained by a reduction in AKT activity in the presence of AβOs (Supplemental 5), which in turn increases lysosomal positioning of TSC and thereby suppresses mTORC1 kinase activity (Norambuena et al., 2017). As expected, AβO treatment increased SOD1 activity in HEK 293 cells (Supplemental Figure 4B) and mouse neuron cultures (Figure 7A and Supplemental Figure 2D and 4C). In addition, an AβO-mediated increase in SOD1 activity was not observed in tau KO neurons (Supplemental 2D). Again, tau expression in Tau KO neurons reduced SOD1 activity by ∼40% and restored the ability of AβOs to increase SOD1 activity (Supplemental 2D). Finally, we explored SOD1 activity in AD human brain. As shown in Figure 7B, SOD1 activity was evaluated by using In-gel activity assays in brain samples taken from a 8 controls and 6 AD donors, SOD1 activity was ∼40% higher in AD compared to normal donors.

**Figure 7.**
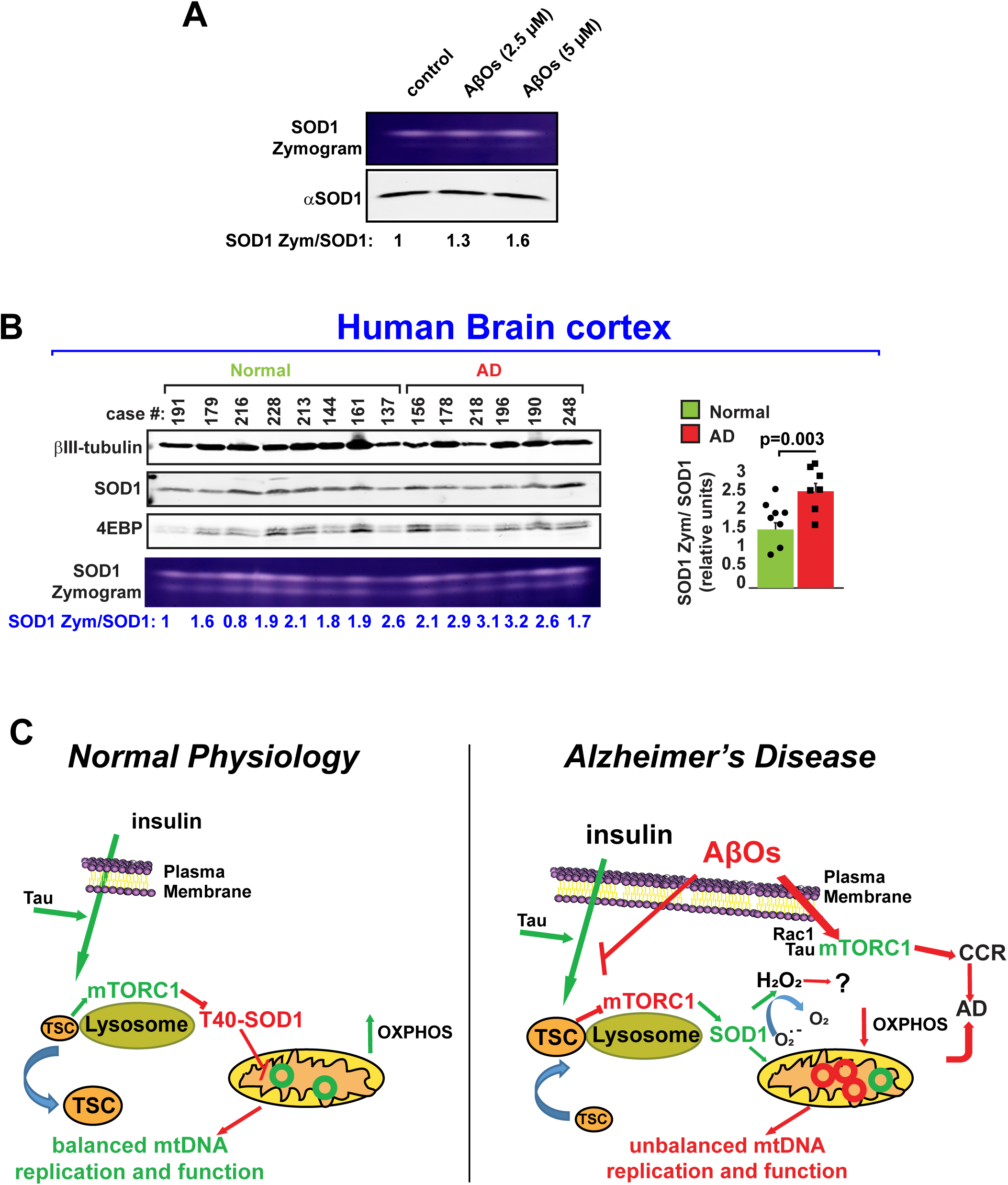
SOD1 activity is increased in AD. **A** SOD1 activity was followed in WT mouse neurons cultures treated with different concentrations of AβO during 1 hour. In-gel activity assays showed that AβO increased SOD1 activity in a dose-dependent manner. Data are representative of three independent assays. **B** SOD1 activity is higher in AD human brains. SOD1 activity assays were analyzed in human brain cell extracts taken from deceased patients affected by AD and compared to normal controls. Samples were analyzed by In-gel activity assays. **C** Model. Under physiological conditions, nutrient stimulated and tau-dependent activation of lysosomal mTORC1 leads to phosphorylation of SOD1 at T40, directly regulating mitochondrial oxidative pathways and mtDNA synthesis by a mechanism independent of Ck1γ2, but likely involving substrates sensitive to oxidation by hydrogen peroxide. Insulin and amino acid-regulated mitochondrial functions are blocked by AβOs, which increased the lysosomal positioning of TSC, likely decreasing mTORC1 activity and stimulating it instead at the PM, where mTORC1 kinase activity triggers cell cycle re-entry (CCR), a frequent prelude to cortical neuron death in Alzheimer’s disease (AD). Thus, SOD1 is at the crossroads of regulating mitochondrial activity through the NiMA pathway and mediating its disruption in AD.

## Discussion

Although considerable progress has been made in the past few decades of AD research, therapies proven to delay or prevent symptom onset, or slow disease progression are still lacking. Of particular relevance is the fact that ∼50% of people with type 2 diabetes (T2D) are eventually affected by sporadic AD (Schrijvers *et al*., 2010). A key feature of T2D is multi-organ metabolic failure, which for brain manifests as reduced insulin sensitivity and increased mitochondrial dysfunction in neurons (De Felice and Ferreira, 2014). Thus, understanding the mechanistic details of the insulin signaling pathway in neurons is pivotal to understanding its pathophysiological effects in AD. Using live incorporation of EdU into mitochondrial nucleoids along with metabolic imaging in live cells and mouse brain, we now expand the mechanistic understanding of an inter-organelle signaling pathway that we recently discovered (Norambuena *et al*., 2018). We show here that activation of lysosomal mTORC1 by insulin and amino acids regulates mitochondrial respiration and mtDNA synthesis by a mechanism controlled by tau and mTORC1-catalyzed phosphorylation of SOD1 at T40. Previous work have reported increased SOD activity in fibroblasts cells lines stablished from AD and normal patients (Zemlan, Thienhaus and Bosmann, 1989) or no changes in brains from control subjects and AD patients (Marklund *et al*., 1985). Using In-gel activity assays, which allows measuring SOD1 activity independently of SOD2 (Weydert and Cullen, 2010), we found higher SOD1 activity in AD human brains, which is also triggered by AβOs in a tau-dependent manner in neuronal cultures, providing additional evidence for how disruption of this lysosome-to-mitochondria signaling pathway represents a seminal step in at least two pathological contexts, AD and tuberous sclerosis, and possibly in ALS as well.

mTOR plays a major physiological role regulating cellular metabolism (Saxton and Sabatini, 2017). These functions require the precise regulation of multiple cellular pathways through serine and/or threonine phosphorylation of potentially >100 substrates (Hsu *et al*., 2011b; Yu *et al*., 2011). Our discovery of the NiMA pathway not only demonstrates that mitochondria quickly react to changes in mTORC1 activity by a mechanism independent of protein synthesis and gene expression, but also established a novel mTORC1 function regulated by its recently discovered substrate, SOD1 (Tsang *et al*., 2018), independently of other well-known mTORC1 substrates, like Bcl-xl, S6 kinase and 4EBP/eIF4E (Norambuena *et al*., 2018). SOD1 is the main regulator of cytosolic redox states (Miao and St. Clair, 2009), but its functional role for mitochondria is not fully understood. We found that SOD1 is part of the neuronal nutrient sensing machinery that mechanistically connects nutrient-mediated activation of lysosomal mTORC1 to mtDNA synthesis and activity. Our observation that inhibition of SOD1 activity with ATN-224 reduced baseline OXPHOS in cells and mouse brains, suggests that interventions targeting SOD1 activity could be beneficial by ameliorating oxidative damage and propagation of oxidized or mutated mtDNA in AD. A note of caution however should be taken considering ATN-224’s pro-apoptotic and pro-oxidant effects in the treatment of cancer (Glasauer *et al*., 2014).

We recently provided mechanistic evidence for how the AβO-tau axis pathologically orchestrates a signaling pathway that at least partially explains how brain insulin resistance may lead to AD by promoting ectopic neuronal cell cycle re-entry and mitochondrial dysfunction (Norambuena *et al*., 2017, 2018; Polanco and Götz, 2018). AβOs can induce an insulin resistant-like phenotype in neurons by sequestering insulin receptors from dendrites and reducing neuronal response to insulin (Zhao *et al*., 2007). AKT phosphorylation at both T308 and S473 leads to full activation (Yang *et al*., 2015), and we found that AβO treatment reduces the phosphate content on AKT, suggesting that AβOs inhibit AKT activity and its downstream targets (Supplemental 5). In fact, we observed that tau re-expression in tau KO neurons restored the lysosomal distribution of the TSC complex, a direct target of AKT phosphorylation by mTORC1 (Menon *et al*., 2014), its sensitivity to nutrient stimulation, and both mTORC1 and SOD1 activities. In addition, tau re-expression also rescued the ability of AβOs to pathologically increase mtDNA synthesis. This ability of tau both to restore “normal” signaling and to facilitate pathological signaling by AβOs is ironic, but possibly explained by the ability of tau to scaffold signaling pathways that became noxious in the presence of AβO, as for example by the activation mTORC1 (Norambuena *et al*., 2017) and Fyn (Ittner *et al*., 2010) at the PM. This paradoxically dual effect should be considered when designing interventions aimed at reducing tau expression in the context of AD and other tauopathies (DeVos *et al*., 2017). In this sense, it is interesting to note that reduction of mTOR genetic dosage or rapamycin treatment (a cellular phenocopy of tau reduction in neurons) ameliorates cognitive deficits in AD transgenic mice (Caccamo *et al*., 2010, 2014). However, rapamycin’s adverse side effects have precluded its broader use in humans (Li, Kim and Blenis, 2014). Further work aimed at understanding the role of SOD1 on the NiMA pathway is necessary for developing more specific interventions. As discuss below, hydrogen peroxide treatment reduced mtDNA synthesis without affecting mitochondrial OXPHOS (Supplemental 6A and B), suggesting that substrates sensitive to oxidation by peroxide are involved and may represent future therapeutic targets.

How does SOD1 regulate mitochondrial function? SOD1 activity has been shown to be required for repressing respiration in yeast and potentially in human cells (Reddi and Culotta, 2013), a process involving SOD1-mediated interaction and stabilization of casein kinase gamma 2 (Ck1γ2), a respiration repressor (Reddi and Culotta, 2013). However, we could not find evidence of such a regulatory mechanism in HEK293 cells or mouse primary neurons, as neither prolonged (24 hours) treatment with ATN-224 nor exposure to AβOs, conditions that respectively abolish or stimulate SOD1 activity, modified Ck1γ2 expression (Supplemental 4A and B). In line with the latter, we did not observe changes in Ck1γ2 expression in human AD brains, in which SOD1 activity was higher compared to non-AD controls (Supplemental 4C and Figure 7B, respectively). Thus, we conclude that in human cells, mouse neurons and human brain, Ck1γ2 expression is not regulated by SOD1 activity and by extension, does not controls mitochondrial function. However, we did observe that hydrogen peroxide (a by-product of SOD1 activity) treatment was able to reduce mtDNA synthesis (Supplemental 6A), suggesting that nutrient-mediated inhibition of mtDNA synthesis is regulated by one or more substrates sensitive to oxidation by H_2_O_2_ (Burgoyne *et al*., 2012). In this respect, it is interesting to note that CaMKII and PKA, whose activity can be modified by H_2_O_2_ (Burgoyne *et al*., 2012), were previously shown to be dysregulated by AβOs (Seward *et al*., 2013; Norambuena *et al*., 2017). Future work is necessary to understand how nutrient-mediated activation of mTORC1, oxidant sensing machinery and kinases activity are integrated to orchestrate proper inter-organelle responses.

The almost exclusively neuronal expression of tau likely conferred neurons with advantages allowing them to perform critical functions during the human lifespan. These functions are not only related to its role as a regulator of cargo transport along microtubules (Ebneth *et al*., 1998; Trinczek *et al*., 1999; Vershinin *et al*., 2007; Dixit *et al*., 2008; Swanson *et al*., 2017), but as we show here, by allowing neurons to properly respond to insulin and nutrients, and to coordinate extracellular cues to cytosolic redox and mitochondrial functioning. The importance of these observations are exemplified by the fact that tau deletion leads to brain insulin resistance in mice (Marciniak *et al*., 2017). Its downstream consequences are also highlighted in this study by showing that tau sustains nutrient-mediated activity of mTORC1 and SOD1. Thus, tau mechanistically connects four key players in AD: insulin and nutrient signaling, mTORC1 function, SOD1 (redox regulation), and mitochondrial activity (De Felice and Ferreira, 2014).

Thus, the results of this study, which are summarizes in Figure 7C, allow us to propose that the tau-mTORC1-SOD1 signaling pathway mechanistically connects mtDNA synthesis and maintenance to nutrient signaling in neurons, opening the possibility that dysregulation of this signaling pathway contributes to mitochondrial oxidative damage and propagation of mutated mtDNA in neurodegeneration. This process potentially could reach pathological levels by changing the balance between normal and damaged mtDNA.

Finally, it is noteworthy that two of the proteins in the pathway described here are emblematic of neurodegeneration: tau in AD and numerous non-Alzheimer’s tauopathies, and SOD1 in ALS, which is an example of a non-Alzheimer’s tauopathy (Lee, Goedert and Trojanowski, 2001). Based on the evidence presented here, it is reasonable to propose that tau perturbed by AβOs leads to SOD1 dysregulation and by extension, mitochondrial dysfunction in AD. Perhaps agents other than AβOs similarly lead to tau perturbation and these downstream effects in other tauopathies and ALS.

## Supplemental Figures Legend

**Supplemental Figure 1.**
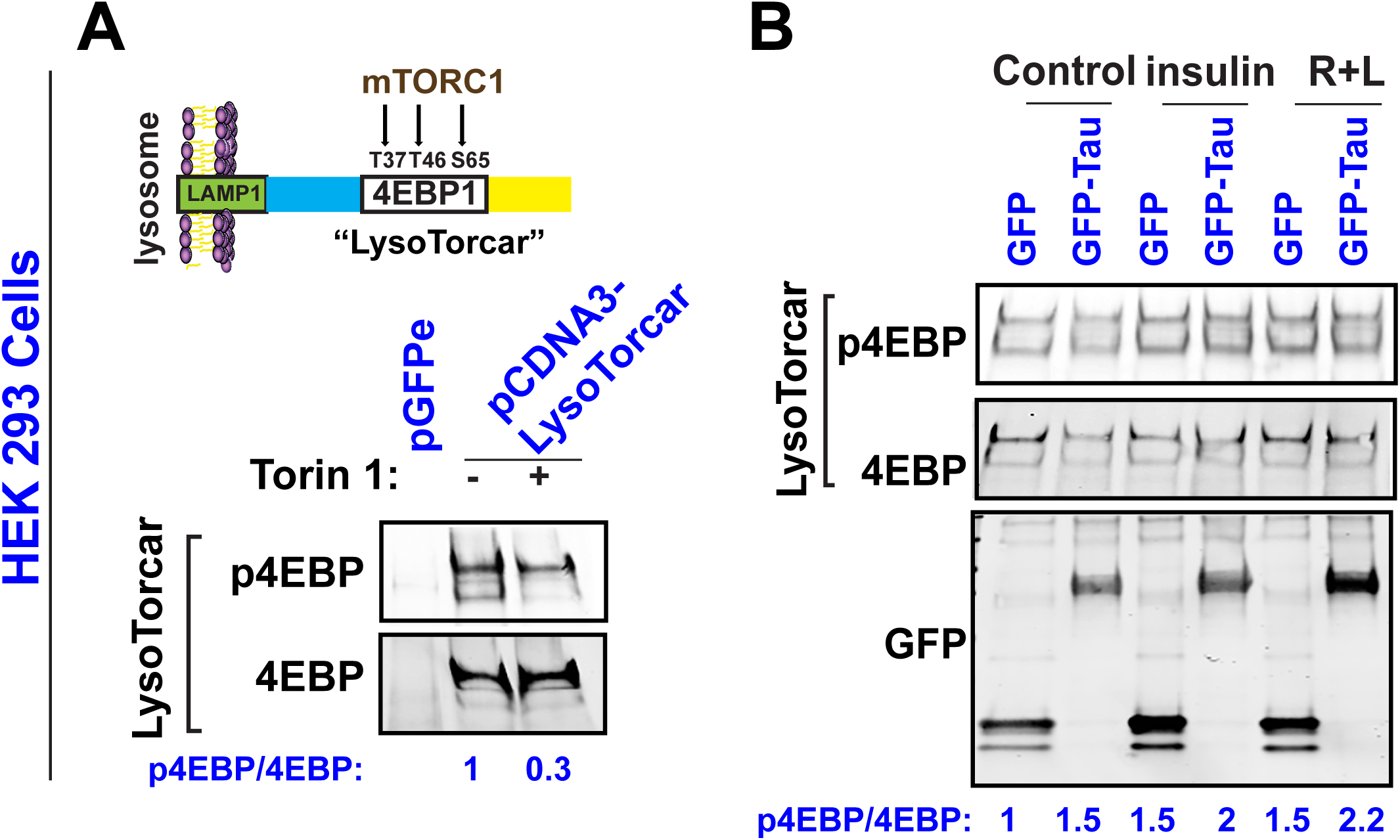
Tau regulates lysosomal activation of mTORC1. **A-B** Lysosomal mTORC1 activity was monitored in HEK 293 cells by following phosphorylation at T37, T46 and S65 in LysoTorcar. Note that expression of GFP-hTau (0N4R) construct increased both base-line and nutrient-stimulated mTORC1 activity. Error bars represent ± s.e.m. Data are representative of two independent assays.

**Supplemental Figure 2.**
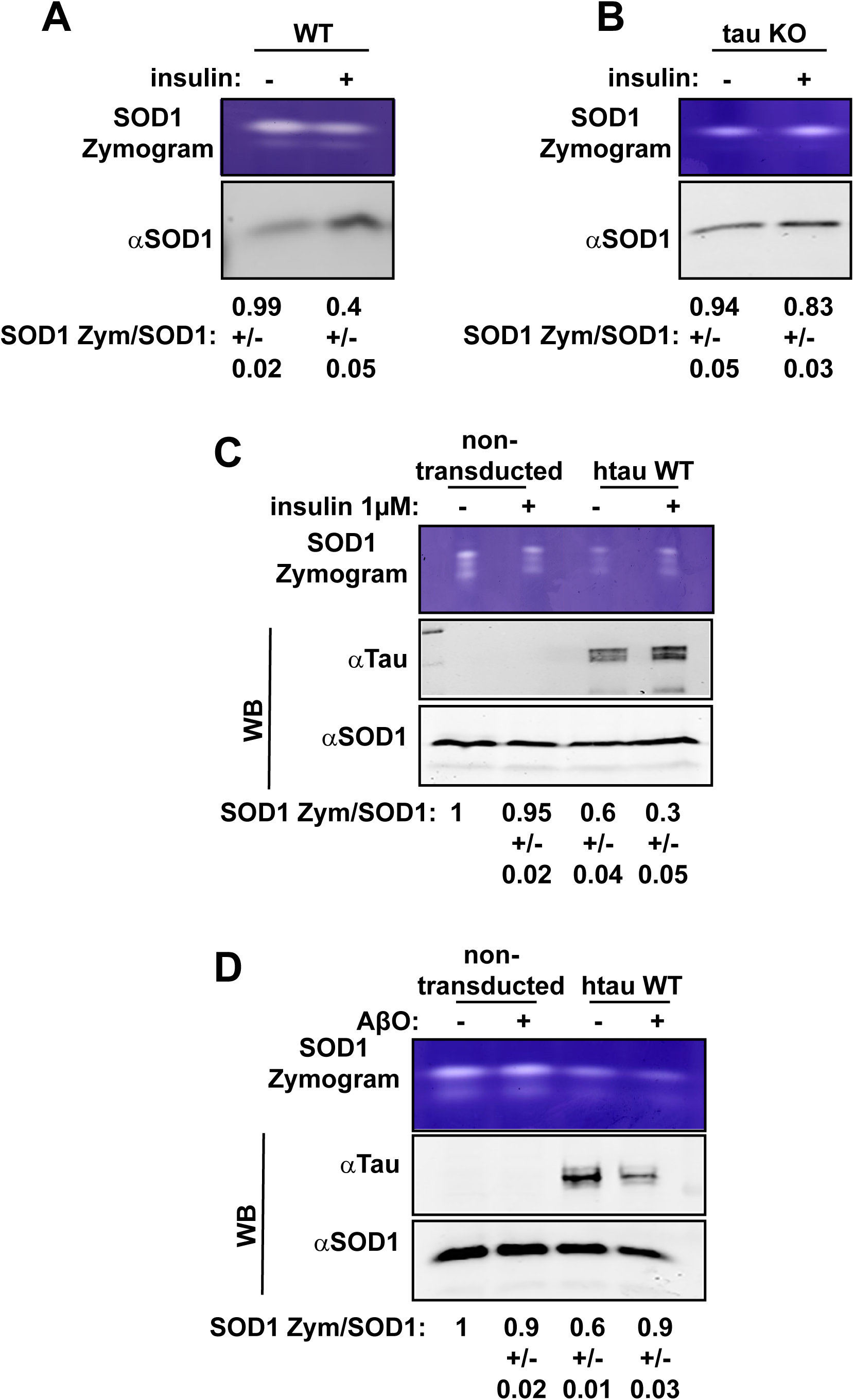
Tau regulates SOD1 activity. **A-B** SOD1 activity was followed in WT (A) or tau KO (B) mouse neurons cultures treated with insulin for 20 minutes. In gel activity assays showed that insulin decreased SOD1 activity by ∼60% and ∼10% when comparing WT and tau KO, respectively. Data are representative of three independent assays. **C** SOD1 activity was followed in unmodified tau KO mouse neurons or after expressing human WT Tau. Neurons were treated with insulin for 20 minutes. In-gel activity assays showed that tau expression alone reduced SOD1 activity by∼40%, and further decreased its activity after insulin treatment. Data are representative of three independent assays. **D** SOD1 activity was followed in unmodified tau KO mouse neurons or after expressing human WT Tau. Neurons were treated with AβOs for 1 hour. In-gel activity assays showed that AβO treatment only increased SOD1 activity in hTau-expressing Tau KO neurons. Data are representative of three independent assays.

**Supplemental Figure 3.**
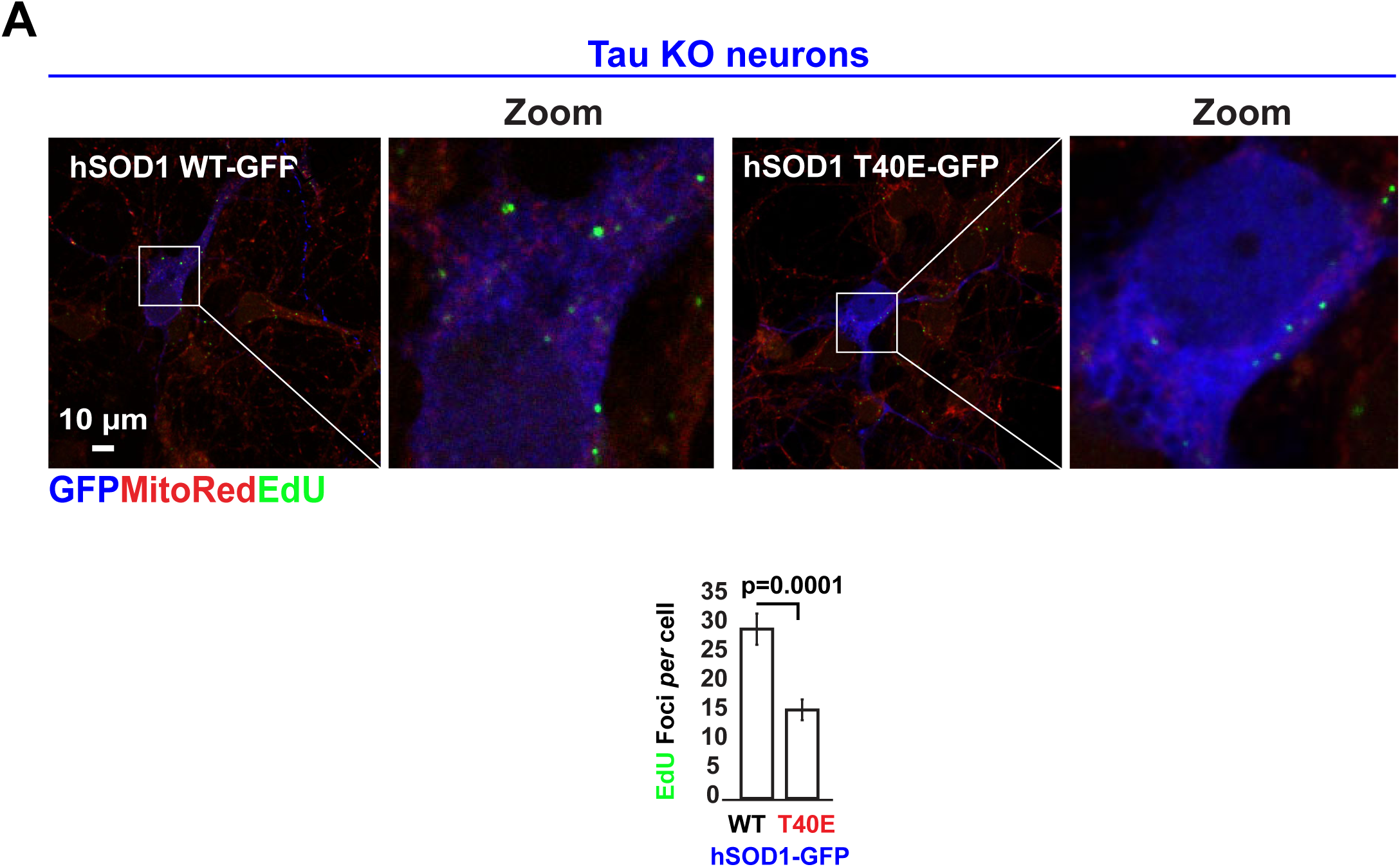
SOD1 phosphorylation in T40 regulates mtDNA synthesis in tau KO neurons. **A** DNA replication in mitochondrial nucleoids was detected in Tau KO mouse cortical neurons expressing either hSOD1^WT^-GFP or hSOD1^T40E^-GFP (29.1±2.6 vs 15.2±1.8 EdU foci/cell, respectively). Each data point in the graph represent the average number of EdU foci/GFP-expressing cell/FoV recorded in this experiment and each FoV contained between 1-2 cells. Error bars represent ± s.e.m. Data are representative of two independent assays.

**Supplemental Figure 4.**
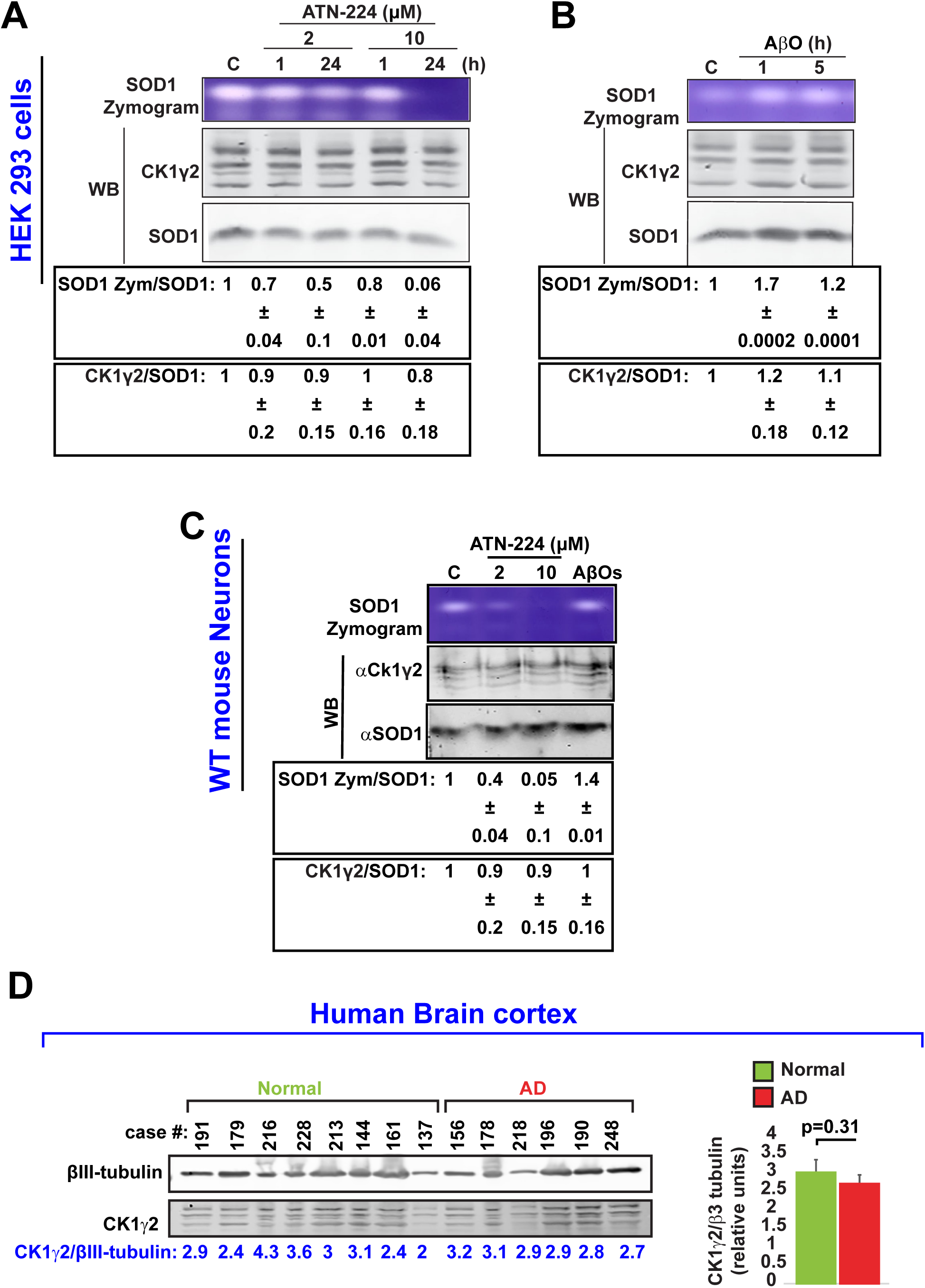
Ck1γ2 expression is independent of SOD1. **A-B** Ck1γ2 expression was evaluated in HEK 293 cells after exposing them during 1 or 24 hours to ATN-224 (2 or 10 µM) or AβOs (3 µM) for 1 or 5 hours. After treatments, cell extracts were prepared for In-gel activity assays or SDS-PAGE and western blots. Note that neither ATN-224 nor AβOs, which respectively decrease or increase SOD1 activity, affected Ck1γ2 expression. Data are representative of three independent assays. **C** Ck1γ2 expression was evaluated in WT mouse neurons after exposing them for 24 hours to ATN-224 (2 or 10 µM) or AβOs (3 µM) for 5 hours. After treatments, cell extracts were prepared for In-gel activity assays or SDS-PAGE and western blots. Similar to results shown in A-B, neither ATN-224 nor AβO affected Ck1γ2 expression in mouse neurons. Data are representative of two independent assays. **D** Ck1γ2 expression was evaluated in human brain. Ck1γ2 expression was assayed by western blots in human brain cell extracts prepared from samples taken from deceased patients affected by AD and compared to normal control subjects.

**Supplemental Figure 5.**
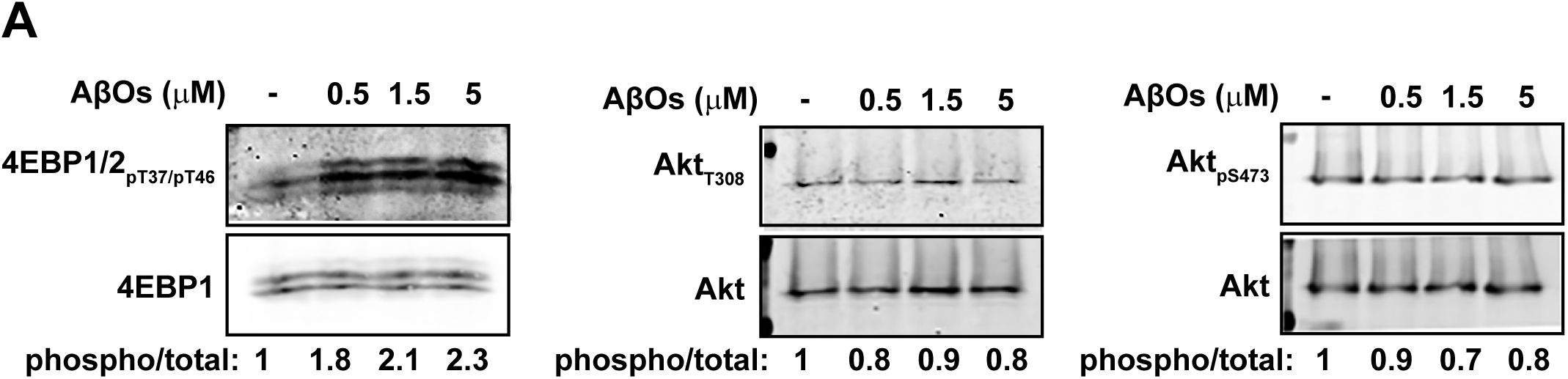
AβO partially reduces AKT phosphorylation. **A** WT cortical neurons were serum-starved in Hank’s balanced salt solution for 2 hours and then treated with AβOs (3 µM) for 30 minutes. After treatments, cell extracts were prepared for SDS-PAGE and western blots against the indicated proteins. AβO-treatment strongly increased phosphorylation of the mTORC1 substrate 4EBP1 (Norambuena *et al*., 2017), concomitantly decreasing AKT phosphorylation in both, PDK1 and mTORC2 sites (Yang *et al*., 2015).

**Supplemental Figure 6.**
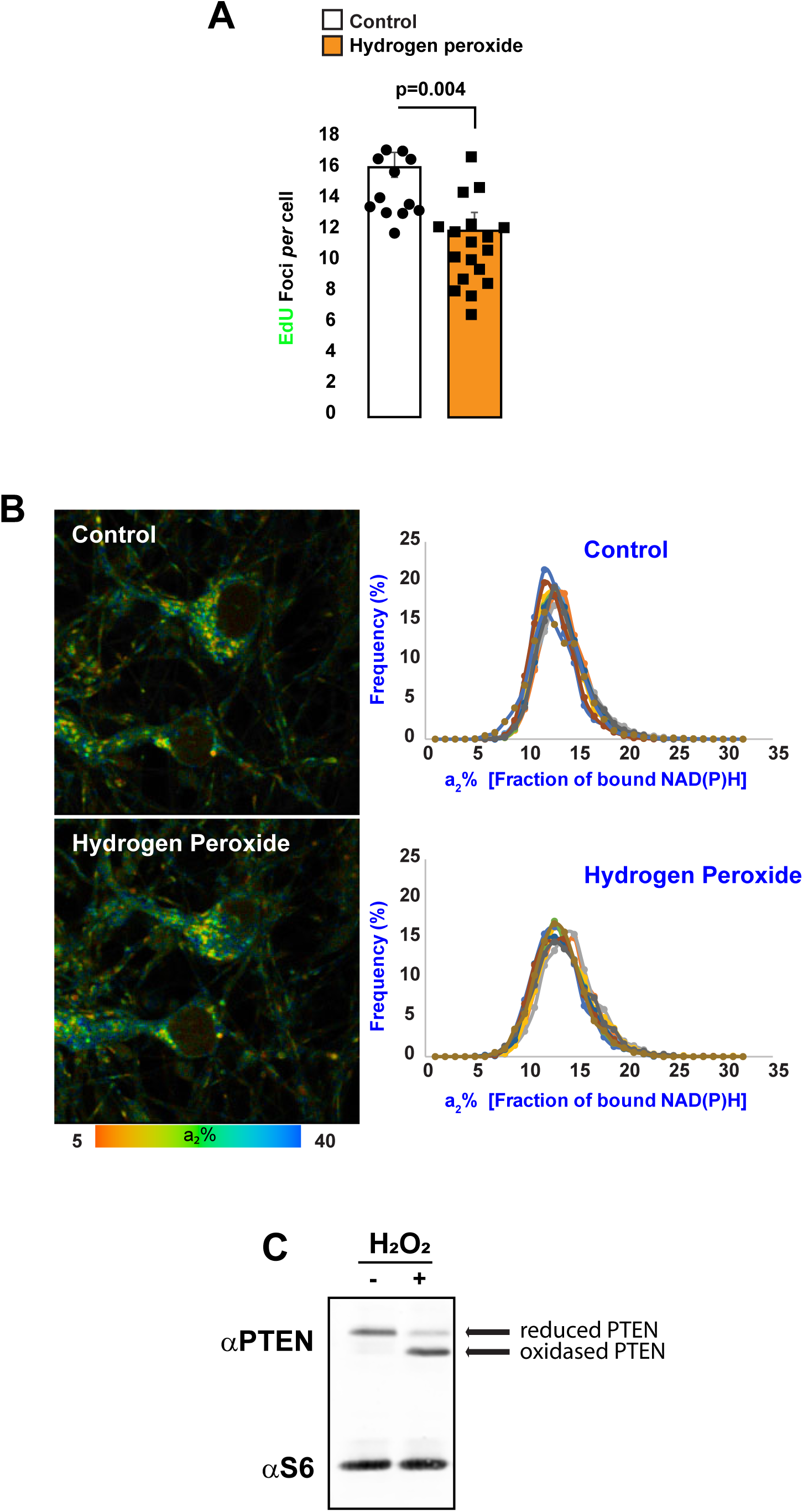
Hydrogen peroxide regulates mtDNA synthesis independently of OXPHOS. **A** DNA replication in mitochondrial nucleoids was detected in WT mouse cortical neurons. Cells were serum-starved in Hank’s balanced salt solution for 2 hours, and then were pulse-labeled for 3 hours with EdU in the absence or presence of hydrogen peroxide (0.5 μM). Quantification of EdU uptake into nucleoids showed that hydrogen peroxide inhibited mtDNA replication by ∼30%. Each data point in the graph represents the average number of EdU foci/cell/FoV recorded in this experiment and each FoV contained between 2-15 cells. Error bars represent ± s.e.m. Data are representative of three independent assays. **B-C** a2% values (starved, and after hydrogen peroxide treatment) were recorded pixel by pixel for WT mouse neurons (B). Each colored line in the upper panel graphs refers to a single field of view containing ∼10 cells, and each colored solid lines in the lower (same color) refers to the same field of view before and after H_2_O_2_ treatment, respectively. Statistical analyses were performed using Student’s two-tailed unpaired t-tests. Under these experimental conditions, ∼70% of PTEN was oxidized by hydrogen peroxide.

**Table EV1.**

| *Antigen* | *Host* | *Application* | *Dilution* | *Source/Cat no.* |
| --- | --- | --- | --- | --- |
| Tau (clone 5) | Mouse | IF - WB | 1/500-1/5000 | LI Binder/<br>Northwestern U |
| Tau (clone R1) | Rabbit | WB | 1/10000 | LI Binder/<br>Northwestern U |
| TSC2 | Rabbit | IF | 1/100 | Cell Signaling<br>Technologies/D93F12 |
| mTOR | Rabbit | IF | 1/100 | Cell Signaling<br>Technologies/2872 |
| Akt(40D4) | Mouse | WB | 1/3000 | Cell Signaling<br>Technologies/2920 |
| Akt (S473) | Rabbit | WB | 1/3000 | Cell Signaling<br>Technologies/9271 |
| Akt (T308) | Rabbit | WB | 1/3000 | Cell Signaling<br>Technologies/9271 |
| 4EBP1 | Rabbit | WB | 1/5000 | Cell Signaling<br>Technologies/55H11 |
| Phospho4EBP1/2<br>(T37/T46) | Rabbit | WB | 1/3000 | Cell Signaling<br>Technologies/9459 |
| Phospho4EBP1/2<br>(S65) | Rabbit | WB | 1/3000 | Cell Signaling<br>Technologies/9451 |
| LAMP1 | Rat | IF | 1/250 | DSHB/1D4B |
| MAP2 | Chicken | IF | 1/10000 | abcam/ab5392 |
| GFP | Mouse | IF-WB | 1/200-1/10000 | NeuroMab UC<br>Davis/N86/8 |
| SOD1 | Rabbit | WB | 1/3000 | Cell Signaling<br>Technologies/37385 |
| SOD1 | Mouse | WB | 1/3000 | Santa Cruz<br>Biotechnology/sc-17767 |
| Casein Kinase 1γ2 | Rabbit | WB | 1/3000 | Invitrogen/PA5-97637 |
| PTEN(138G6) | Rabbit | WB | 1/2000 | Cell Signaling<br>Technologies/9559 |
| S6 ribosomal<br>protein(5G10) | Rabbit | WB | 1/2000 | Cell Signaling<br>Technologies/2217 |
| AlexaFluor 488 | Goat Anti<br>Chicken IgG | IF | 1/500 | Life<br>technologies/A11039 |
| AlexaFluor 488 | Goat Anti<br>Rabbit IgG | IF | 1/500 | Life<br>technologies/A11008 |
| AlexaFluor 488 | Goat Anti<br>Mouse IgG | IF | 1/500 | Life<br>technologies/A11001 |
| AlexaFluor 594 | Goat Anti<br>Rabbit IgG | IF | 1/500 | Life<br>technologies/A11012 |
| AlexaFluor 594 | Goat Anti<br>Mouse IgG | IF | 1/500 | Life<br>technologies/A11005 |
| AlexaFluor 647 | Goat Anti<br>Rat IgG | IF | 1/500 | Life<br>technologies/A21247 |
| IRDye 800 | Goat Anti<br>Mouse IgG | WB | 1/10000 | LICOR/926-32210 |
| IRDye 680 | Goat Anti<br>Rabbit IgG | WB | 1/10000 | LICOR/936-68171 |

## Materials and Methods

### Cell lines

Human embryonic kidney (HEK293) were grown in DMEM/F12 media (GIBCO) supplemented with HyClone cosmic calf serum (GE Healthcare) and 50 µg/ml gentamycin (GIBCO).

Human tuberous sclerosis fibroblasts with a polymorphisms in *TSC1* gene (catalog number GM06149 and GM06121) were obtained from the Coriell Institute, and were cultured according to the vendor’s specifications in Eagle’s minimal essential medium (GIBCO) supplemented with 15% Optima fetal bovine serum (Atlanta Biologicals).

### Cultured mouse neurons

Brain cortices were dissected from E17/18 WT (C57/Bl6) or tau knockout mice (Dawson *et al*., 2001), and primary neuron cultures were prepared as described previously (Norambuena *et al*., 2017, 2018). Experiments were performed using cultures grown for at least 10 days in Neurobasal medium (GIBCO) supplemented with B27 (GIBCO) and 50 µg/ml gentamycin before being used for experiments, except that lentivirus transductions were performed 3 days in advance of all other experimental procedures.

### Amyloid-β oligomers (AβOs)

Lyophilized, synthetic Aβ_1–42_ (AnaSpec) was dissolved in 1,1,1,3,3,3-hexafluoro-2-propanol (Sigma-Aldrich) to ∼1 mM and evaporated overnight at room temperature. The dried powder was resuspended for 5 minutes at room temperature in 40-50 µl dimethylsulfoxide to ∼1 mM and sonicated for 10 minutes in a water bath. To prepare oligomers, the dissolved, monomeric peptide was diluted to ∼400 µl (100 µM final concentration) in Neurobasal medium (GIBCO), incubated 24-48 hours at 4° C with rocking, and then centrifuged at 14,000 g for 15 minutes to remove fibrils. For all experiments, AβOs were diluted into tissue culture medium to a final concentration of ∼1.5 µM total Aβ_1–42._

### Lentivirus production and infection

Lentiviral particles for shRNA knockdowns and hTau 2N4R and 0N4R protein expression in mouse cortical neurons were prepared as follows. The pBOB-NepX or pLKO.1 expression plasmids, and the packaging vectors, pSPAX2 and pMD2.G (Addgene plasmids 12260 and 12259, respectively) were transfected using Lipofectamine 3000 (ThermoFisher) into HEK293T cells grown in 15 cm Petri dishes to ∼80% confluence in DMEM (GIBCO) supplemented with 10% HyClone cosmic calf serum (GE Healthcare). Each transfection was with 15 µg total DNA at a 50%/37.5%/12.5% ratio of expression vector/pSPAX2/pMD2.G. Lentivirus-conditioned medium was collected 24 and 48 hours after the start of transfection. Lentiviral particles were concentrated in a Beckman Coulter Optima LE-80K ultracentrifuge for 2 hours at 23,000 rpm (95,000 g_av_) at 4° C in an SW28 rotor, resuspended in 400 µl Neurobasal medium and stored at −80° C in 20 µl aliquots. Cultured neurons were transduced in Neurobasal/B27 medium and incubated for 72 hours before assays were performed.

### Human Brain Cell Extracts

This study was approved by the Institutional Review Board for Health Sciences Research at the University of Virginia. Human frontal cortex brain samples were previously acquired via generous donation from the University of Virginia Brain Resource Facility. A total of 8 patients free from pathological evidence of Alzheimer’s disease (mean age 73.5±3.9 years, 50% female, average post-mortem interval (PMI) 9.1±1.6 hours) were compared to 6 patients with Alzheimer’s pathology (mean age 76.8±2.3 years, 50% female, average PMI 7.5±1.8 hours). There was no significant difference in age (p=0.53) or PMI (p=0.51) between normal and AD subjects. Portioned brain samples in 1.5ml Eppendorf tubes were thawed on ice, and one scoop of 0.5mm zirconium oxide beads was added along with 100 – 200 μL of lysis buffer to cover sample and beads. Lysis buffer consisted of RIPA buffer (Bioworld, #42020024-2); 1:100 HALT™ protease inhibitor cocktail and 1:100 EDTA (Thermo Scientific, #78430); 1:100 phosphatase inhibitor cocktail 2 (Sigma Aldrich, #P5726); and 1:100 phosphatase inhibitor cocktail 3 (Sigma Aldrich, #P0044). This mixture was placed for 5-15 minutes in a bullet blender at 4°C until fully homogenized, after which homogenate was placed in a centrifuge at 10,000 RPM for 10 minutes at 4°C. Finally, supernatant was transferred to a new tube and used as protein lysate for subsequent SOD1 activity assays (see below) and western blots.

### Superoxide Dismutase 1 Activity Assays

Superoxide dismutase 1 activity was evaluated as described previously (Weydert and Cullen, 2010). This method detects SOD activity from freshly as well as frozen specimens (Weydert and Cullen, 2010). Briefly, mammalian cells were washed three times in phosphate-buffered saline (Fisher Scientific BP665), lysed in 50 mM phosphatase buffer (0.05M KH_2_PO_4_, 0.05M K_2_HPO_4_, pH 7.8) with a Sonicator (FisherScientific FB120) set at 40% power for 30s on ice. Cell lysates were diluted in native 2X sample buffer (Bio Rad 1610738). Protein samples were separated in 12% native PAGE gels, which were stained with nitro blue tetrazolium chloride (Abcam, catalog number ab146262), as described previously (Weydert and Cullen, 2010). To visualize SOD1 and SOD2 activities, gels were rinsed with water three times and placed on a light box for 20 minutes to 2 hours. SOD1 and SOD2 bands were identified either by sodium cyanide treatment in the gel staining step (Weydert and Cullen, 2010) or shRNA mediated knockdown of SOD1 in mouse neurons.

### Oxygen Consumption Assay

WT mouse cortical neurons grown in 96 well plates (black plate, clear bottom, Costar catalog number 3603) were serum-starved in Hank’s balanced salt solution, as described earlier (Norambuena *et al*., 2018), then treated for 1 hour with the SOD1 inhibitor and copper chelator, ATN-224 Cayman Chemical (cat no. 23553). Oxygen consumption was assayed using the Extracellular O_2_ Consumption Assay (Abcam catalog number ab197243).

### PTEN Oxidation assay

Hydrogen peroxide-induced oxidation of PTEN was performed as described by Juarez et al. (Juarez *et al*., 2008) with few modifications. Briefly, WT neurons were serum starved in HBSS during 2 hours, then treated with 500nM hydrogen peroxide for 15 min. Neurons were washed in cold PBS on ice, lysed in RIPA buffer (1% Nonidet P40, 0.25% sodium deoxicolate,150mM NaCl, 50mM Tris-HCl pH=7.5, 200μM sodium orthovanadate, 10μM NaF, 100nM okadaic acid) supplemented with 10mM N-ethylmaleimide (a kind gift by Dr. Anthony Spano. Department of Biology, University of Virginia) during 30 minutes at 4°C. Samples were run on a nonreducing SDS-PAGE and transferred to nitrocellulose membranes (BioRad cat. No. 1620112). The same membranes were incubated with antibodies against PTEN and S6 ribosomal protein as a loading control. Oxidized PTEN runs faster than the reduced form on a SDS-PAGE (Juarez *et al*., 2008).

### cDNA Constructs and shRNA Sequences

Mouse TSC2 knockdowns was described by us before (Norambuena *et al*., 2017). pcDNA3 LysoTorcar was a kind gift of Dr. Jin Zhang (University of California, San Diego). Human 2N4R tau vector (Norambuena *et al*., 2017) was amplified by PCR and transferred to pBOB-NepX between the AgeI and HpaI sites using the following primers: forward 5’ TTA ACC GGT ATG GCT GAG CCC CGC CAG 3’ and reverse 5’ ATT GTTAAC CTA CAA ACC CTG CTT GGC CAG 3’. The same strategy was use to transfer hTau 0N4R WT (see below) to the pBOB NepX lentiviral vector. To knockdown TSC2, a shTSC2-pLKO.1 construct was described previously (Norambuena *et al*., 2017)

The human SOD1 phospho-null (T40A) and phospho-mimetic (T40E) constructs were generated using the NEB Q5^®^ Site-Directed Mutagenesis Kit (cat noE0554S) using as a template the human SOD1 construct pF146 pSOD1WTAcGFP1described below. Primers design and procedure was done following manufacturer recommendations (http://nebasechanger.neb.com/).

The following constructs were from Addgene:

1) pRK7-FLAG-TSC1 (plasmid 8995; gift from Dr. John Blenis).
2) pRK5-GFP-hTAU 0N4R WT (plasmid 46904; gift from Dr. Karen Ashe).
3) pF146 pSOD1WTAcGFP1 (plasmid 26407: gift from Elizabeth Fisher)
4) pSPAX2 and pMD2.G (plasmids 12259 and 12260; gift from Didier Trono)

The following plasmids were purchased from The RNAi Consortium of the Broad Institute;

1) Mouse Nprl3 (TRCN0000175195)
2) Mouse SOD1 (TRCN0000101046)

### Immunoblotting

Samples were resolved by SDS-PAGE using either 10 or 12% acrylamide/bis-acrylamide gels and transferred to 0.22 µm nitrocellulose (Bio-Rad). Membranes were blocked with Odyssey blocking buffer (LI-COR Biosciences), and were incubated with primary antibodies and secondary IRDye-labeled antibodies (see Extended Data Table 1) diluted into antibody buffer (Odyssey blocking buffer diluted 1:1 in PBS/0.1% Tween 20). All antibody incubations were for 1 hour at room temperature or overnight at 4° C, and 3 washes of 5 minutes each with PBS/0.1% Tween 20 were performed after each antibody step. Membranes were dried between sheets of filter paper prior to quantitative imaging with an Odyssey imaging station (LI-COR Biosciences).

### Antibodies

See table 1.

### NADH and NADPH measurement

#### Time-correlated single photon counting (TCSPC) Fluorescence Lifetime Imaging Microscopy (FLIM)

FLIM was recorded on a Zeiss LSM-780 NLO confocal/multiphoton microscopy system comprising an inverted Axio Observer Z1 microscope, an X-Cite 120PC Q mercury arc light source (Excelitas Technologies) for cell selection, a motorized stage for automated scanning, an IR Chameleon Vision-II ultrafast Ti:sapphire laser for multiphoton excitation (Coherent), a Zeiss 40X 1.3 NA oil immersion Planapo objective, an environmental chamber (PeCon GmbH, Germany) that envelops the microscope stage to control temperature and CO_2_ level, and a 3-channel FLIM system based on three HPM-100-40 GaAsP-based hybrid detectors and 3 SPC-150 TCSPC boards (Becker & Hickl). The SPC-150 boards are synchronized with the 2-photon excitation laser and the Zeiss LSM-780 NLO scan head signal. Ex 740nm; Em450/50 nm.

#### Imaging

Mouse neurons, and HEK293 cells were grown in 35-mm glass-bottom dishes, and maintained at 37° C in 5% CO_2_/95% air on the stage of the Zeiss LSM-780 NLO microscope. All cultures were serum-starved in Hank’s balanced salt solution (HBSS) for 2 hours before addition of ATN 224 to 2 μM, and imaged before and 60 minutes after treatment. The laser was tuned to 740 nm with an average power of 7 mW at the specimen plane, and NAD(P)H fluorescence was collected using a 450-500 nm emission filter (Objective lens, 20x). For each experiment, 5–10 field of views were recorded in the descanned mode, and then each field of view was subjected to a 40-second acquisition in the non-descanned mode. The laser power and acquisition time were selected to ensure enough photons per pixel while avoiding photodamage to cells. Next, ROIs corresponding to mitochondria were selected from the NAD(P)H photon image for lifetime analysis. Both the lifetimes and fractions of free and enzyme-bound NAD(P)H were calculated on a per pixel basis, from which pseudo-color images of the bound NAD(P)H and histograms of the frequency distribution of a_2_% (fraction of bound NAD(P)H) were generated.

#### Processing

FLIM images were processed with SPCImage software (v5; Becker & Hickl) with non-linear least square for optimal fitting. Data images were exported for analysis using SPCImage to establish the free and enzyme-bound fractions of NAD(P)H. The intensity images were used to document mitochondrial morphology and to generate ROIs with a custom plug-in for ImageJ (http://imagej.net/Welcome) to capture the discrete and heterogeneous nature of mitochondrial dynamics. A custom ImageJ macro extracted parameters of interest, most notably the fraction of enzyme-bound NAD(P)H. A custom macro in Microsoft Excel further processed the data by individual ROI and produced histograms, charts and statistics, followed by merging the different field of views for charting.

#### Co-localization assays

As we had done earlier (Norambuena *et al*., 2017), co-localization of mTOR with LAMP1 or Mitotracker CMXRos was quantified by an ImageJ plug-in (http://rsbweb.nih.gov/ij/index.html) for the Manders coefficient, as described before which is based on the Pearson Correlation Coefficient (Dunn, Kamocka and McDonald, 2011).

### Other fluorescence microscopy procedures

#### Primary mouse neuron transfections

6-7 days old cultures of WT or Tau KO neurons growing on no. 1.5 thickness, 12 mm round glass coverslips in 24-well culture dishes were transfected with human SOD1-GFP WT, TA or TE using Lipofectamine 2000 (ThermoFisher). Each transfection was performed by diluting 0.5 µg of plasmid in 50 µls of plain Neurobasal medium. On a different tube, 1µl of Lipofectamine 2000 (ThermoFisher) was diluted in 50 µls of the same medium. The final DNA/Lipofectamine complexes (100uls) were added to the cells maintained in 100 µls of conditioned medium. 5-6 hours after transfection, 300 µls of previously saved conditioned medium was added back to the cultures. EdU uptake assay was performed 24 hours later as described below. Cells were fixed with 3.7% para-formaldehyde in PBS, and the coverslips were sealed to glass slides using Fluormount G (ThermoFisher).

#### Human tuberous sclerosis fibroblasts

Cells growing on #1.5 thickness, 12 mm round glass coverslips in 24-well dishes were transfected with 0.5 µg of pRK7-FLAG-TSC1 using Lipofectamine 2000 (ThermoFisher) following the same procedure described in the previous paragraph. Expression of exogenously encoded TSC1 was monitored 24 hours later by immunofluorescence. Cells were rinsed in PBS and fixed for 15 minutes in 3.7% para-formaldehyde. Next, they were washed and permeabilized in washing buffer (0.2% Tween 20 in PBS) 3 times for 5 minutes each and incubated for 60 minutes in blocking buffer (PBS version; LI-COR Biosciences). Fixed cells were then incubated for 1 hour each with primary and secondary antibodies diluted into blocking buffer supplemented with Tween 20 to 0.2%, with several PBS washes after each antibody step. Finally, coverslips were mounted onto glass slides using Fluormount-G (ThermoFisher). Samples were imaged on a Nikon Eclipse Ti equipped with a Yokogawa CSU-X1 spinning disk head, 60X 1.4 NA Planapo objective, a Hamamastu Flash 4.0 scientific CMOS camera, and 405 nm, 488 nm, 561 nm and 640 nm lasers.

#### Mitochondrial DNA (mtDNA) replication experiments

To visualize mtDNA replication by we used Click-it chemistry as described by others (Lewis, Uchiyama and Nunnari, 2016). EdU is a thymidine analog that readily incorporates into active replicating DNA, which can be detected by a copper catalyzed reaction that covalently links (“clicks”) the alkyne group in EdU to the picolyl azide group in an Alexa Fluor® dye (Salic and Mitchison, 2008). Mitochondria were labeled with Mitotracker CMXRos (Invitrogen catalog number M7512), and AlexaFluor 647-EdU-tagged nucleoids were revealed using the Click-It chemistry (Salic and Mitchison, 2008). Briefly, WT mouse cortical neurons grown on glass coverslips were serum-starved in HBBS for 2 hours in the presence or absence of AβOs (see below) and aphidicolin (7.5µM) before cells were treated for 3 hours with nutrients (1 µM insulin plus 0.8 mM L-leucine [Sigma Aldrich catalog numbers I5500 and L8000, respectively] and 0.4mM L-arginine [Acros Organics catalog number 105001000)]). For immunofluorescence, cells were incubated with 100 nM Mitotracker Red CMXRos for 60 minutes before being assayed. The immunofluorescence labeling procedure was as described for human tuberous sclerosis fibroblasts. Mitochondrial nucleoids were counted using the particle analysis plugging for ImageJ (https://imagej.net/Particle_Analysis).

### Multiparametric Photoacoustic Microscopy (MP-PAM)

12-16 week-old male CD-1 mice were used in this study. All animal procedures were approved by the Institutional Animal Care and Use Committee at Washington University in St. Louis. Following hair removal, a surgical incision was made in the scalp. The exposed skull was cleaned. Then, the mouse skull region over a 3×3 mm^2^ ROI was carefully removed to expose the cortex for topical application of ATN-224. After craniotomy, the anesthetized mouse was transferred to a stereotaxic instrument. The animal body temperature was maintained at 37° C via a heating pad and the local brain temperature was also maintained at 37°C via a temperature-controlled water tank. Ultrasound gel was applied between the open-skull window and water tank for acoustic coupling. Following the baseline imaging, the ultrasound gel was gently removed and a solution of ATN-224 (4µM) was applied topically. The exposed mouse cortex was treated with the SOD1 inhibitor solution for 80 minutes. Then, the ROI was covered again with ultrasound gel and subjected to post-treatment MP-PAM imaging (Cao *et al*., 2017). For 2P-FLIM imaging of the live mouse brain, the same pre-imaging procedure was used, but in this case for 12 week old male C57BL/6 mice. After surgery, the mice were transferred to a stereotaxic frame at the side of the Zeiss LSM-780 microscope. An objective converter was used to convert the inverted LSM-780 to upright configuration. A 20x/NA 0.85 water immersion objective lens was coupled to the converter and gently moved to the cranial window for FLIM imaging.

### Statistics

For 2P-FLIM assays and statistical analyses were performed using Student’s two-tailed unpaired t-test for experiments with a single treatment unless otherwise stated. Since there were thousands of ROIs obtained per image, the average of the ROIs for each field of view was calculated to reduce the sample size and thus the quantity of false positives. All data obtained follow the assumptions of normal distributions. All data displayed follow a frequency distribution relative to a_2_% bound to NADPH.

The paired t-test was used for analyzing bar graphs shown in Figs: 1B, D and F, 2C-F, 3 A and B, 4B, D, F and G, 5B, 6A-C, and 7B.

For MP-PAM assays, vessel segmentations and quantitative analysis were done following our established protocol(Cao *et al*., 2017). The paired t-test was used for comparing the cerebral hemodynamics and oxygen metabolism before and after application of the compounds, which is shown in Figure 5E.

## Acknowledgments

We are grateful for financial support from NIH/NIA (grant R01AG067048 to AN) the Alzheimer’s and Related Diseases Research Award Fund (grant 17-5 to AN), University of Virginia’s Brain Institute and Virginia Alzheimer’s Disease Center Leadership Award (AP, AN and GSB), the Owens Family Foundation (GSB), NIH/NIA (grant RF1 AG051085 to GSB); NIH/Office of the Director for funds to purchase a Zeiss 780 microscope that was used in this study (OD016446 to AP), the Cure Alzheimer’s Fund (GSB), the Alzheimer’s Association (grants ZEN-16-363266 to GSB), Webb and Tate Wilson (GSB), The Virginia Chapter of the Lady’s Auxiliary of the Fraternal Order of Eagles (GSB), and the University of Virginia President’s Fund for Excellence (GSB).

## Author contributions

AN conceived, designed and initiated the study, performed most of the experiments, analyzed all data and co-wrote the paper. GSB analyzed data and co-wrote the paper. RC performed 2P-FLIM experiments in the live mouse brain. HW analyzed 2P-FLIM experiments in the live cells and mouse brains. AP provided extensive technical support for 2P-FLIM. XS performed experiments in tuberous sclerosis human fibroblasts. EP performed Ck1γ2 experiments, DBW and NS prepared mouse neuron cultures. NS and SH performed and analyzed experiments involving MP-PAM. LP and HF provided cell extracts from human brain samples. All authors read, edited and approved submission of the manuscript.

## Conflict of Interests

The authors have no conflicts of interest to report.

## References

Adav, S. S., Park, J. E. and Sze, S. K. (2019) ‘Quantitative profiling brain proteomes revealed mitochondrial dysfunction in Alzheimer’s disease’, Molecular Brain. doi: 10.1186/s13041-019-0430-y.

Arendt, T. et al. (2010) ‘Selective cell death of hyperploid neurons in Alzheimer’s disease.’, The American journal of pathology, 177(1), pp. 15–20. doi: 10.2353/ajpath.2010.090955.

Balaban, R. S., Nemoto, S. and Finkel, T. (2005) ‘Mitochondria, oxidants, and aging.’, Cell, 120(4), pp. 483–95. doi: 10.1016/j.cell.2005.02.001.

Blacker, T. S. et al. (2014) ‘Separating NADH and NADPH fluorescence in live cells and tissues using FLIM’, Nature Communications, 5. doi: 10.1038/ncomms4936.

Blanchard, B. J. et al. (1993) ‘A mitochondrial DNA deletion in normally aging and in alzheimer brain tissue’, NeuroReport. doi: 10.1097/00001756-199306000-00051.

Burgoyne, J. R. et al. (2012) ‘Hydrogen Peroxide Sensing and Signaling by Protein Kinases in the Cardiovascular System’, Antioxidants & Redox Signaling. Mary Ann Liebert, Inc., publishers, 18(9), pp. 1042–1052. doi: 10.1089/ars.2012.4817.

Burté, F. et al. (2014) ‘Disturbed mitochondrial dynamics and neurodegenerative disorders’, Nature Reviews Neurology, 11(1), pp. 11–24. doi: 10.1038/nrneurol.2014.228.

Caccamo, A. et al. (2010) ‘Molecular interplay between mammalian target of rapamycin (mTOR), amyloid-beta, and Tau: effects on cognitive impairments.’, The Journal of biological chemistry, 285(17), pp. 13107–20. doi: 10.1074/jbc.M110.100420.

Caccamo, A. et al. (2014) ‘Genetic reduction of mammalian target of rapamycin ameliorates Alzheimer’s disease-like cognitive and pathological deficits by restoring hippocampal gene expression signature.’, The Journal of neuroscience : the official journal of the Society for Neuroscience, 34(23), pp. 7988–98. doi: 10.1523/JNEUROSCI.0777-14.2014.

Calkins, M. J. et al. (2011) ‘Impaired mitochondrial biogenesis, defective axonal transport of mitochondria, abnormal mitochondrial dynamics and synaptic degeneration in a mouse model of Alzheimer’s disease’, Human Molecular Genetics, 20(23), pp. 4515–4529. doi: 10.1093/hmg/ddr381.

Camandola, S. and Mattson, M. P. (2017) ‘Brain metabolism in health, aging, and neurodegeneration’, The EMBO Journal. doi: 10.15252/embj.201695810.

Cao, R. et al. (2017) ‘Functional and oxygen-metabolic photoacoustic microscopy of the awake mouse brain’, NeuroImage, 150, pp. 77–87. doi: 10.1016/j.neuroimage.2017.01.049.

Carrieri, G. et al. (2001) ‘Mitochondrial DNA haplogroups and APOE4 allele are non-independent variables in sporadic Alzheimer’s disease’, Human Genetics. doi: 10.1007/s004390100463.

Chagnon, P. et al. (1999) ‘Phylogenetic analysis of the mitochondrial genome indicates significant differences between patients with Alzheimer disease and controls in a French-Canadian founder population’, American Journal of Medical Genetics. doi: 10.1002/(SICI)1096-8628(19990702)85:1<20::AID-AJMG6>3.0.CO;2-K.

Che, M. et al. (2016) ‘Expanding roles of superoxide dismutases in cell regulation and cancer’, Drug Discovery Today. doi: 10.1016/j.drudis.2015.10.001.

Chen, H. and Chan, D. C. (2009) ‘Mitochondrial dynamics--fusion, fission, movement, and mitophagy--in neurodegenerative diseases.’, Human molecular genetics, 18(R2), pp. R169–76. doi: 10.1093/hmg/ddp326.

Chen, Y. et al. (2016) ‘Mitochondrial DNA Rearrangement Spectrum in Brain Tissue of Alzheimer’s Disease: Analysis of 13 Cases’, PLoS ONE. doi: 10.1371/journal.pone.0154582.

Chidambaram, M. V., Barnes, G. and Frieden, E. (1984) ‘Inhibition of ceruloplasmin and other copper oxidases by thiomolybdate’, Journal of Inorganic Biochemistry. Elsevier, 22(4), pp. 231–239. doi: 10.1016/0162-0134(84)85008-4.

Collins, J.-A. et al. (2015) ‘Relating oxygen partial pressure, saturation and content: the haemoglobin-oxygen dissociation curve.’, *Breathe (Sheffield*, England), 11(3), pp. 194–201. doi: 10.1183/20734735.001415.

Corral-Debrinski, M. et al. (1994) ‘Marked changes in mitochondrial dna deletion levels in alzheimer brains’, Genomics. doi: 10.1006/geno.1994.1525.

Coskun, P. E., Beal, M. F. and Wallace, D. C. (2004) ‘Alzheimer’s brains harbor somatic mtDNA control-region mutations that suppress mitochondrial transcription and replication’, Proceedings of the National Academy of Sciences of the United States of America. doi: 10.1073/pnas.0403649101.

Cottrell, D. A. et al. (2001) ‘Mitochondrial enzyme-deficient hippocampal neurons and choroidal cells in AD’, Neurology. doi: 10.1212/WNL.57.2.260.

Crane, P. K. et al. (2013) ‘Glucose Levels and Risk of Dementia’, New England Journal of Medicine. Massachusetts Medical Society, 369(6), pp. 540–548. doi: 10.1056/NEJMoa1215740.

Croteau, E. et al. (2018) ‘A cross-sectional comparison of brain glucose and ketone metabolism in cognitively healthy older adults, mild cognitive impairment and early Alzheimer’s disease’, Experimental Gerontology. doi: 10.1016/j.exger.2017.07.004.

Dawson, H. N. et al. (2001) ‘Inhibition of neuronal maturation in primary hippocampal neurons from τ deficient mice’, Journal of Cell Science, 114(6), pp. 1179–1187. doi: 10.1242/jcs.114.6.1179.

DeVos, S. L. et al. (2017) ‘Tau reduction prevents neuronal loss and reverses pathological tau deposition and seeding in mice with tauopathy’, Science Translational Medicine, 9(374). doi: 10.1126/scitranslmed.aag0481.

Dixit, R. et al. (2008) ‘Differential regulation of dynein and kinesin motor proteins by tau’, Science, 319(5866). doi: 10.1126/science.1152993.

DuBoff, B., Feany, M. and Götz, J. (2013) ‘Why size matters - balancing mitochondrial dynamics in Alzheimer’s disease’, Trends in Neurosciences, pp. 325–335. doi: 10.1016/j.tins.2013.03.002.

Dunn, K. W., Kamocka, M. M. and McDonald, J. H. (2011) ‘A practical guide to evaluating colocalization in biological microscopy’, AJP: Cell Physiology, 300(4), pp. C723–C742. doi: 10.1152/ajpcell.00462.2010.

Ebneth, A. et al. (1998) ‘Overexpression of tau protein inhibits kinesin-dependent trafficking of vesicles, mitochondria, and endoplasmic reticulum: Implications for Alzheimer’s disease’, Journal of Cell Biology, 143(3). doi: 10.1083/jcb.143.3.777.

De Felice, F. G. and Ferreira, S. T. (2014) ‘Inflammation, defective insulin signaling, and mitochondrial dysfunction as common molecular denominators connecting type 2 diabetes to Alzheimer Disease’, Diabetes, pp. 2262–2272. doi: 10.2337/db13-1954.

Gao, X. et al. (2002) ‘Tsc tumour suppressor proteins antagonize amino-acid-TOR signalling’, Nature Cell Biology. doi: 10.1038/ncb847.

Glasauer, A. et al. (2014) ‘Targeting SOD1 reduces experimental non–small-cell lung cancer’, The Journal of Clinical Investigation. The American Society for Clinical Investigation, 124(1), pp. 117–128. doi: 10.1172/JCI71714.

Gordon, B. A. et al. (2018) ‘Spatial patterns of neuroimaging biomarker change in individuals from families with autosomal dominant Alzheimer’s disease: a longitudinal study’, The Lancet Neurology. doi: 10.1016/S1474-4422(18)30028-0.

Guo, X. et al. (2017) ‘Amyloid β-induced redistribution of transcriptional factor EB and lysosomal dysfunction in primary microglial cells’, Frontiers in Aging Neuroscience. doi: 10.3389/fnagi.2017.00228.

Gustafsson, C. M., Falkenberg, M. and Larsson, N.-G. (2016) ‘Maintenance and Expression of Mammalian Mitochondrial DNA’, Annual Review of Biochemistry. doi: 10.1146/annurev-biochem-060815-014402.

Hsu, P. P. et al. (2011a) ‘The mTOR-regulated phosphoproteome reveals a mechanism of mTORC1-mediated inhibition of growth factor signaling.’, *Science (New York*, N.Y.), 332(6035), pp. 1317–22. doi: 10.1126/science.1199498.

Hsu, P. P. et al. (2011b) ‘The mTOR-regulated phosphoproteome reveals a mechanism of mTORC1-mediated inhibition of growth factor signaling.’, *Science (New York*, N.Y.), 332(6035), pp. 1317–22. doi: 10.1126/science.1199498.

Hudson, G. et al. (2012) ‘No consistent evidence for association between mtDNA variants and Alzheimer disease’, Neurology. doi: 10.1212/WNL.0b013e31824e8f1d.

Inczedy-Farkas, G. et al. (2014) ‘Mitochondrial DNA mutations and cognition: A case-series report’, Archives of Clinical Neuropsychology. doi: 10.1093/arclin/acu016.

Ishii, K. et al. (1996) ‘Decreased medial temporal oxygen metabolism in Alzheimer’s disease shown by PET.’, *Journal of nuclear medicine : official publication*, Society of Nuclear Medicine, 37(7), pp. 1159–65. Available at: http://europepmc.org/abstract/MED/8965188 (Accessed: 25 July 2014).

Ittner, L. M. et al. (2010) ‘Dendritic function of tau mediates amyloid-β toxicity in alzheimer’s disease mouse models’, Cell, 142(3). doi: 10.1016/j.cell.2010.06.036.

Juarez, J. C. et al. (2008) ‘Superoxide dismutase 1 (SOD1) is essential for H2O2-mediated oxidation and inactivation of phosphatases in growth factor signaling.’, Proceedings of the National Academy of Sciences of the United States of America, 105(20), pp. 7147–7152. doi: 10.1073/pnas.0709451105.

Kapogiannis, D. and Mattson, M. P. (2011) ‘Disrupted energy metabolism and neuronal circuit dysfunction in cognitive impairment and Alzheimer’s disease.’, The Lancet. Neurology, 10(2), pp. 187–98. doi: 10.1016/S1474-4422(10)70277-5.

Labbé, K., Murley, A. and Nunnari, J. (2014) ‘Determinants and functions of mitochondrial behavior.’, Annual review of cell and developmental biology, 30, pp. 357–91. doi: 10.1146/annurev-cellbio-101011-155756.

Lakatos, A. et al. (2010) ‘Association between mitochondrial DNA variations and Alzheimer’s disease in the ADNI cohort’, Neurobiology of Aging. doi: 10.1016/j.neurobiolaging.2010.04.031.

Lakowicz, J. R. (2006) *Principles of Fluorescence Spectroscopy Principles of Fluorescence Spectroscopy*, *Principles of fluorescence spectroscopy*, Springer, New York, USA, 3rd edn, 2006. doi: 10.1007/978-0-387-46312-4.

Lee, V. M.-Y., Goedert, M. and Trojanowski, J. Q. (2001) ‘Neurodegenerative Tauopathies’, Annual Review of Neuroscience. Annual Reviews, 24(1), pp. 1121–1159. doi: 10.1146/annurev.neuro.24.1.1121.

Lewis, S. C., Uchiyama, L. F. and Nunnari, J. (2016) ‘ER-mitochondria contacts couple mtDNA synthesis with Mitochondrial division in human cells’, Science, 353(6296), p. aaf5549. doi: 10.1126/science.aaf5549.

Li, J., Kim, S. G. and Blenis, J. (2014) ‘Rapamycin: One drug, many effects’, Cell Metabolism. doi: 10.1016/j.cmet.2014.01.001.

Liang, W. S. et al. (2008) ‘Alzheimer’s disease is associated with reduced expression of energy metabolism genes in posterior cingulate neurons.’, Proceedings of the National Academy of Sciences of the United States of America, 105(11), pp. 4441–6. doi: 10.1073/pnas.0709259105.

Manczak, M., Calkins, M. J. and Reddy, P. H. (2011) ‘Impaired mitochondrial dynamics and abnormal interaction of amyloid beta with mitochondrial protein Drp1 in neurons from patients with Alzheimer’s disease: Implications for neuronal damage’, Human Molecular Genetics, 20(13), pp. 2495–2509. doi: 10.1093/hmg/ddr139.

Marciniak, E. et al. (2017) ‘Tau deletion promotes brain insulin resistance’, Journal of Experimental Medicine, 214(8). doi: 10.1084/jem.20161731.

Marklund, S. L. et al. (1985) ‘Superoxide dismutase isoenzymes in normal brains and in brains from patients with dementia of Alzheimer type’, Journal of the Neurological Sciences, 67(3), pp. 319–325. doi: https://doi.org/10.1016/0022-510X(85)90156-X.

Menon, S. et al. (2014) ‘Spatial control of the TSC complex integrates insulin and nutrient regulation of mtorc1 at the lysosome’, Cell, 156(4), pp. 1771–1785. doi: 10.1016/j.cell.2013.11.049.

Miao, L. and St. Clair, D. K. (2009) ‘Regulation of superoxide dismutase genes: Implications in disease’, Free Radical Biology and Medicine. doi: 10.1016/j.freeradbiomed.2009.05.018.

Mosconi, L. et al. (2007) ‘Maternal family history of Alzheimer’s disease predisposes to reduced brain glucose metabolism’, Proceedings of the National Academy of Sciences, 104(48), pp. 19067–19072. doi: 10.1073/pnas.0705036104.

Murakami, K. et al. (2011) ‘SOD1 (copper/zinc superoxide dismutase) deficiency drives amyloid β protein oligomerization and memory loss in mouse model of Alzheimer disease.’, The Journal of biological chemistry, 286(52), pp. 44557–44568. doi: 10.1074/jbc.M111.279208.

Nair, P. M. and Mason, H. S. (1966) ‘Reconstitution of cytochrome c oxidase from an apo-enzyme and Cu(I)’, Biochemical and Biophysical Research Communications. doi: 10.1016/0006-291X(66)90261-0.

Ning, B. et al. (2015) ‘Ultrasound-aided Multi-parametric Photoacoustic Microscopy of the Mouse Brain.’, Scientific reports. Nature Publishing Group, 5, p. 18775. doi: 10.1038/srep18775.

Norambuena, A. et al. (2017) ‘mTOR and neuronal cell cycle reentry: How impaired brain insulin signaling promotes Alzheimer’s disease’, Alzheimer’s and Dementia, 13(2), pp. 152–167. doi: 10.1016/j.jalz.2016.08.015.

Norambuena, A. et al. (2018) ‘A novel lysosome-to-mitochondria signaling pathway disrupted by amyloid-β oligomers.’, The EMBO journal. EMBO Press, p. e100241. doi: 10.15252/embj.2018100241.

Nunnari, J. and Suomalainen, A. (2012) ‘Mitochondria: In sickness and in health’, Cell, pp. 1145–1159. doi: 10.1016/j.cell.2012.02.035.

Orlova, K. A. and Crino, P. B. (2010) ‘The tuberous sclerosis complex’, Annals of the New York Academy of Sciences, 1184(1), pp. 87–105. doi: 10.1111/j.1749-6632.2009.05117.x.

Polanco, J. C. et al. (2018) ‘Amyloid-β and tau complexity - Towards improved biomarkers and targeted therapies’, Nature Reviews Neurology. doi: 10.1038/nrneurol.2017.162.

Polanco, J. C. and Götz, J. (2018) ‘ Are you TORCing tau me? Amyloid-β blocks the conversation between lysosomes and mitochondria ’, The EMBO Journal. doi: 10.15252/embj.2018100839.

Prather, P. and de Vries, P. J. (2004) ‘Behavioral and cognitive aspects of tuberous sclerosis complex.’, Journal of child neurology, 19(9), pp. 666–74. doi: 10.1177/08830738040190090601.

Reddi, A. R. and Culotta, V. C. (2013) ‘SOD1 integrates signals from oxygen and glucose to repress respiration’, Cell. doi: 10.1016/j.cell.2012.11.046.

Rolfe, D. F. and Brown, G. C. (1997) ‘Cellular energy utilization and molecular origin of standard metabolic rate in mammals’, Physiological Reviews. American Physiological Society, 77(3), pp. 731–758. doi: 10.1152/physrev.1997.77.3.731.

Sakamoto, H. et al. (2018) ‘A family case with germline TSC1 and mtDNA mutations developing bilateral eosinophilic chromophobe renal cell carcinomas without other typical phenotype of tuberous sclerosis’, Journal of Clinical Pathology. BMJ Publishing Group, 71(10), pp. 936–943. doi: 10.1136/jclinpath-2018-205211.

Salic, A. and Mitchison, T. J. (2008) ‘A chemical method for fast and sensitive detection of DNA synthesis in vivo.’, Proceedings of the National Academy of Sciences of the United States of America, 105(7), pp. 2415–2420. doi: 10.1073/pnas.0712168105.

Saxton, R. A. and Sabatini, D. M. (2017) ‘mTOR Signaling in Growth, Metabolism, and Disease’, Cell, pp. 960–976. doi: 10.1016/j.cell.2017.02.004.

Schrijvers, E. M. C. et al. (2010) ‘Insulin metabolism and the risk of Alzheimer disease: The Rotterdam Study’, Neurology. doi: 10.1212/WNL.0b013e3181ffe4f6.

Seward, M. E. M. E. et al. (2013) ‘Amyloid-β signals through tau to drive ectopic neuronal cell cycle re-entry in Alzheimer’s disease.’, Journal of cell science, 126(Pt 5), pp. 1278–86. doi: 10.1242/jcs.1125880.

Shadel, G. S. and Horvath, T. L. (2015) ‘Mitochondrial ROS Signaling in Organismal Homeostasis’, Cell. doi: 10.1016/j.cell.2015.10.001.

Sheng, Z.-H. and Cai, Q. (2012) ‘Mitochondrial transport in neurons: impact on synaptic homeostasis and neurodegeneration.’, Nature reviews Neuroscience, 13(2), pp. 77–93. doi: 10.1038/nrn3156.

Smalley, S. L. (1998) ‘Autism and tuberous sclerosis’, Journal of Autism and Developmental Disorders, pp. 407–414. doi: 10.1023/A:1026052421693.

Suomalainen, A. and Battersby, B. J. (2017) ‘Mitochondrial diseases: the contribution of organelle stress responses to pathology’, Nature Reviews Molecular Cell Biology. doi: 10.1038/nrm.2017.66.

Swanson, E. et al. (2017) ‘Extracellular Tau Oligomers Induce Invasion of Endogenous Tau into the Somatodendritic Compartment and Axonal Transport Dysfunction’, Journal of Alzheimer’s Disease. IOS Press, 58, pp. 803–820. doi: 10.3233/JAD-170168.

Thiele, E. A. (2004) ‘Managing Epilepsy in Tuberous Sclerosis Complex’, Journal of Child Neurology, 19(9), pp. 680–686. doi: 10.1177/08830738040190090801.

Trifunovic, A. et al. (2004) ‘Premature ageing in mice expressing defective mitochondrial DNA polymerase’, Nature, 429(6990), pp. 417–423. doi: 10.1038/nature02517.

Trinczek, B. et al. (1999) ‘Tau regulates the attachment/detachment but not the speed of motors in microtubule-dependent transport of single vesicles and organelles’, Journal of Cell Science, 112(14).

Tsang, C. K. et al. (2018) ‘SOD1 Phosphorylation by mTORC1 Couples Nutrient Sensing and Redox Regulation’, Molecular Cell. doi: 10.1016/j.molcel.2018.03.029.

Valentine, J. S., Doucette, P. A. and Potter, S. Z. (2005) ‘Copper-zinc superoxide dismutase and amyotrophic lateral sclerosis’, Annual Review of Biochemistry. doi: 10.1146/annurev.biochem.72.121801.161647.

Valla, J., Berndt, J. D. and Gonzalez-Lima, F. (2001) ‘Energy hypometabolism in posterior cingulate cortex of Alzheimer’s patients: Superficial laminar cytochrome oxidase associated with disease duration’, Journal of Neuroscience. doi: 10.1523/jneurosci.21-13-04923.2001.

Veal, E. A., Day, A. M. and Morgan, B. A. (2007) ‘Hydrogen Peroxide Sensing and Signaling’, Molecular Cell. doi: 10.1016/j.molcel.2007.03.016.

Vershinin, M. et al. (2007) ‘Multiple-motor based transport and its regulation by Tau’, Proceedings of the National Academy of Sciences of the United States of America, 104(1). doi: 10.1073/pnas.0607919104.

Vyas, S., Zaganjor, E. and Haigis, M. C. (2016) ‘Mitochondria and Cancer’, Cell, pp. 555–566. doi: 10.1016/j.cell.2016.07.002.

Van Der Walt, J. M. et al. (2004) ‘Analysis of European mitochondrial haplogroups with Alzheimer disease risk’, Neuroscience Letters. doi: 10.1016/j.neulet.2004.04.051.

Wang, W. et al. (2020) ‘Mitochondria dysfunction in the pathogenesis of Alzheimer’s disease: Recent advances’, Molecular Neurodegeneration. doi: 10.1186/s13024-020-00376-6.

Weisiger, R. A. and Fridovich -, I. (1973) ‘Mitochondrial superoxide dismutase. Site of synthesis and intramitochondrial localization’, Journal of Biological Chemistry. doi: 10.1016/S0021-9258(19)43735-6.

Weydert, C. J. and Cullen, J. J. (2010) ‘Measurement of superoxide dismutase, catalase and glutathione peroxidase in cultured cells and tissue’, Nature Protocols, 5(1). doi: 10.1038/nprot.2009.197.

Wiedemann, F. R. et al. (2002) ‘Mitochondrial DNA and respiratory chain function in spinal cords of ALS patients’, *Journal of Neurochemistry*. John Wiley & Sons, Ltd, 80(4), pp. 616–625. doi: https://doi.org/10.1046/j.0022-3042.2001.00731.x.

Yang, G. et al. (2015) ‘A Positive Feedback Loop between Akt and mTORC2 via SIN1 Phosphorylation’, Cell Reports. doi: 10.1016/j.celrep.2015.07.016.

Yoon, E. J. et al. (2009) ‘Intracellular amyloid beta interacts with SOD1 and impairs the enzymatic activity of SOD1: implications for the pathogenesis of amyotrophic lateral sclerosis’, Experimental & Molecular Medicine, 41(9), pp. 611–617. doi: 10.3858/emm.2009.41.9.067.

Yu, Y. et al. (2011) ‘Phosphoproteomic analysis identifies Grb10 as an mTORC1 substrate that negatively regulates insulin signaling.’, *Science (New York*, N.Y.), 332(6035), pp. 1322–6. doi: 10.1126/science.1199484.

Zemlan, F. P., Thienhaus, O. J. and Bosmann, H. B. (1989) ‘Superoxide dismutase activity in Alzheimer’s disease: possible mechanism for paired helical filament formation.’, Brain research. Netherlands, 476(1), pp. 160–162. doi: 10.1016/0006-8993(89)91550-3.

Zhao, W.-Q. et al. (2007) ‘Amyloid beta oligomers induce impairment of neuronal insulin receptors’, The FASEB Journal, 22(1), pp. 246–260. doi: 10.1096/fj.06-7703com.

Zhong, Z. et al. (2018) ‘New mitochondrial DNA synthesis enables NLRP3 inflammasome activation’, Nature. doi: 10.1038/s41586-018-0372-z.

Zhou, X. et al. (2015) ‘Dynamic Visualization of mTORC1 Activity in Living Cells’, Cell Reports. doi: 10.1016/j.celrep.2015.02.031.

